# Mucosal tissue NK cells directly mediate tissue protection and repair during infection

**DOI:** 10.1101/2025.04.04.647286

**Authors:** Sarah C. Vick, Eva Domenjo-Vila, Marie Frutoso, Raisa A. Glabman, Lakshmi S. Warrier, Sean M. Hughes, Anna C. Kirby, Michael F. Fialkow, Florian Hladik, Martin Prlic, Jennifer M. Lund

**Author notes:** UMR1227, LBAI, University Brest, Inserm, Brest, France.

## Abstract

Preserving barrier integrity while mounting effective immunity is essential at mucosal surfaces. In examining the immune cells that mediate both inflammation and tissue homeostasis, we uncovered a dual role for natural killer (NK) cells in barrier immunity. While NK cells are known to control viral infections, here we identify a previously unrecognized reparative function for NK cells in mucosal tissues. Using single-cell RNA sequencing and high-dimensional flow cytometry, we reveal a distinct population of human mucosal NK cells marked by tissue residency, immunoregulatory profiles, and limited cytotoxic potential at homeostasis, yet highly responsive to inflammatory cues. In a mouse model of acute HSV-2 infection, NK cells are required for limiting infection-associated tissue damage. Mucosal NK cells express the epithelial growth factor amphiregulin (Areg), and their depletion leads to increased tissue barrier damage despite preserved viral clearance, suggesting a novel role in tissue protection. Mechanistically, we demonstrate that the barrier-derived cytokines IL-18 and IL-33 induce Areg expression by both human and mouse NK cells, linking local inflammatory cues to reparative NK cell programming that is able to potentiate wound healing. Together, our findings reveal a context-dependent, dual function of mucosal NK cells in immune defense and mucosal tissue protection, expanding current models of NK cell biology.

## Introduction

Mucosal surfaces are particularly vulnerable to infection as they directly interface with the environment and so are frequently exposed to pathogens. Thus, mucosal tissues have unique immune mechanisms to limit pathogen replication while still maintaining tissue integrity. Immune populations important for direct cell killing and promoting inflammation, including CD8+ T cells and natural killer (NK) cells, work together to eliminate infections^1, 2^. NK cells detect virally infected cells through a combination of activating and inhibitory receptors^3^, while T cells largely depend on antigen recognition to kill infected cells or pathogens. These cells both act through the secretion of pro-inflammatory cytokines like IFNγ and direct killing of cells with molecules such as perforin and granzyme B. While the immune responses critical to protect the host from pathogens at barrier sites are gaining attention, less is known about the resolution of inflammation after infection and the return to homeostasis. Immunoregulation at mucosal sites is important as excess inflammation can lead to tissue damage and thus compromised primary tissue function, so we sought to understand cellular mechanisms critical for tissue healing and homeostasis in mucosal sites in the context of infection.

NK cells are present in the circulation and within tissues. Tissue-resident NK cells have been described in multiple lymphoid and non-lymphoid sites, including the small intestines^4^, lungs^5^, salivary glands^6^, liver^7, 8, 9^, uterus,^10, 11^ and skin^12^. Tissue-resident NK cells differ from circulating NK cells in several ways, including expression of tissue retention markers^4^ and reduced cytotoxicity^13, 14^. It is yet to be fully elucidated if tissue NK cells are transiently recruited from peripheral blood during inflammation^12, 15^ or if tissue NK cells are developmentally distinct from circulating NK cells and constitute a long-lived, self-renewing tissue-resident population^4, 5, 7, 16, 17^. However, regardless of their ontogeny, the unique signatures of NK cells in mucosal and barrier tissue sites suggest unique functional capacities that require a thorough investigation in individual anatomic sites and disease contexts.

While characterization of NK cells in the lower female genital mucosa is incomplete, it is known that NK cells play critical roles during host defense against sexually transmitted viral infections, such as human immunodeficiency virus (HIV)^18, 19^ and herpes simplex virus 2 (HSV-2). During primary HSV-2 infection in mice, NK cells provide a crucial early source of IFNγ ^20, 21^, and depletion of NK cells leads to increased viral burden in vaginal tissue and the spinal cord, demonstrating the critical role of NK cells in controlling infection^22^.

While pro-inflammatory cytokines, such as IFNγ, are crucial for pathogen clearance, we hypothesize that unrestrained NK cell activity at mucosal sites can drive chronic inflammation and collateral tissue damage. This raises the possibility that mucosal NK cells act as a double-edged sword during viral infection, requiring a balance between pathogen control and barrier maintenance for optimal host health. Here, we characterize NK cells from human and murine vaginal tissue and blood, revealing that vaginal NK cells exhibit a unique transcriptional and protein signature. Vaginal NK cells are largely quiescent at homeostasis yet poised for robust responses to inflammatory cues. Strikingly, these mucosal NK cells also produce the tissue repair factor amphiregulin, and their activity supports epithelial barrier integrity and tissue repair following HSV-2 infection. Together, our findings uncover a novel mechanism by which mucosal NK cells mediate an active balance between anti-pathogen effector function and direct preservation and repair of host tissue health.

## Results

### scRNA-seq analysis reveals that vaginal NK cells express tissue repair gene signatures

We performed single-cell RNA sequencing (scRNA-seq) on vaginal tissue (VT) and paired peripheral blood (PBMC) samples enriched for CD45+ immune cells from 4 healthy participants undergoing reconstructive vaginal surgery (Figure 1A). We analyzed the scRNA-seq data and calculated principal component analysis and Uniform Manifold Approximation and Projection (UMAP) for dimensionality reduction. We used SingleR to guide cell type recognition, and manual cell annotation was confirmed by examining the top 8 differentially expressed genes across all clusters (Extended Data Figure 1A). We obtained sequences from approximately 32,000 cells and found most immune cell subsets represented in all donors, including B cells, T cells, dendritic cells, monocytes, and natural killer (NK) cells (Figure 1B, Extended Data Figure 1B-C). CD8 and CD4 Effector/Memory T cells made up most of the immune cells from the VT and PBMC samples, while regulatory T cells, B cells, and NK cells also made up a substantial proportion of the immune cells from each donor (Extended Data Figure 1B). Immune cells from the VT and PBMC clustered separately (Figure 1C), suggesting that these populations may have different gene signatures, consistent with our previous work focused on vaginal tissue T cells ^23, 24, 25^, and from published work showing that NK cells from tissues, including the lung and liver, are distinct from circulating or lymphoid cells.^4, 5, 6, 7, 8, 9, 10, 11, 12, 13, 14^

**Figure 1.**
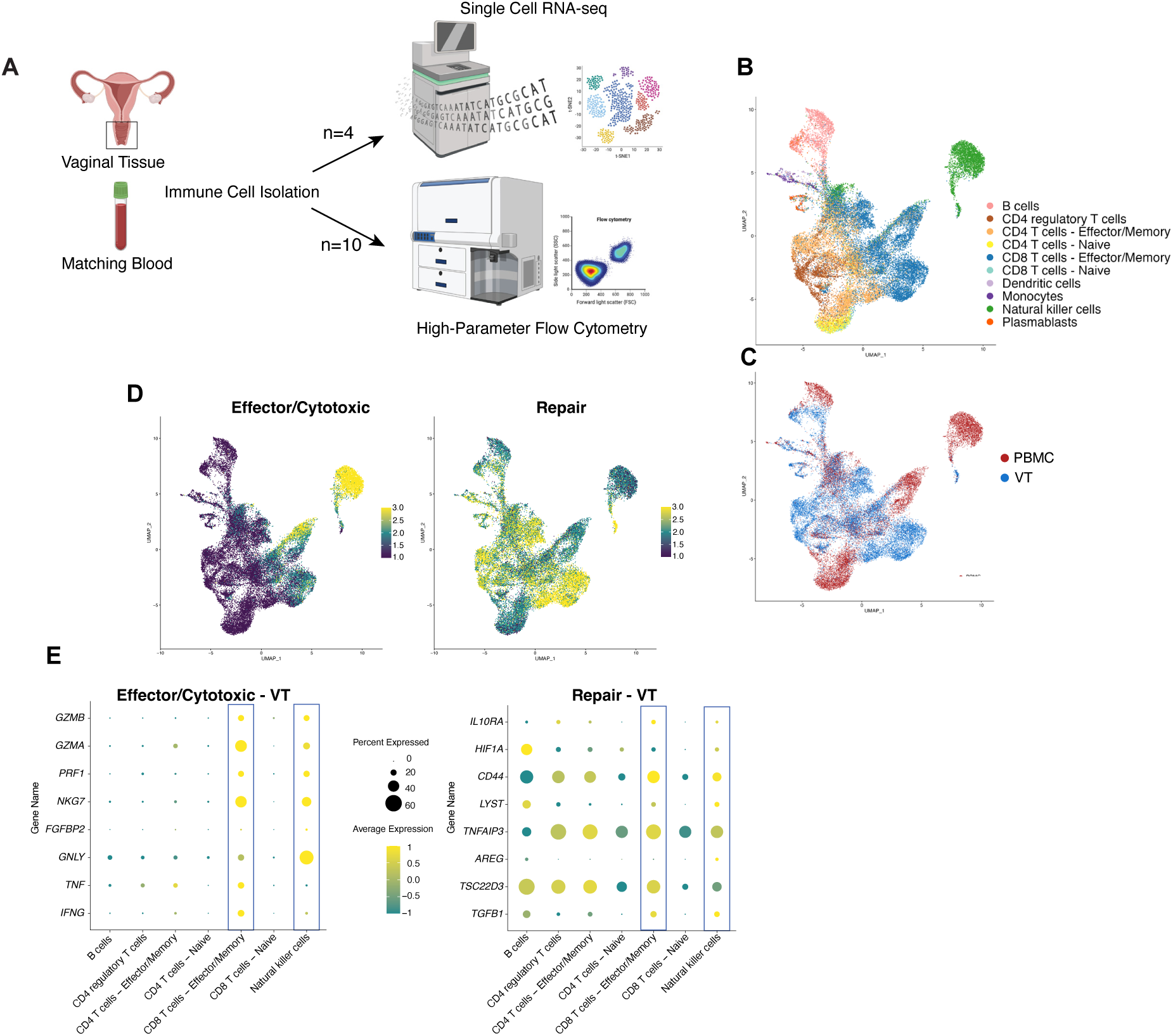
scRNA-seq analysis reveals vaginal tissue (VT) immune cell populations with repair gene signature. We performed single-cell RNA sequencing (scRNA-seq) on vaginal tissue (VT) and matching blood (PBMC) samples enriched for CD45+ immune cells from 4 healthy donors. Fastq files were processed via the standard Rhapsody analysis pipeline on Seven Bridges. After read filtering, reads were aligned to a reference genome and annotated, barcodes and UMIs were counted, followed by determining putative cells. The final output was analyzed in R using Seurat. Principal component analysis was performed to calculate UMAP and graph-based clustering with a resolution between 0.2 and 0.6. For cell annotation, we applied SingleR. **A**) Experimental design for human studies.**B**) UMAP plot colored by clustering based on transcripts. **C**) UMAP plot based on tissue type. **D**) UMAP plot showing enrichment of effector/cytotoxic or repair gene signature. **E**) Scaled dot plot showing average transcript expression (color) and percent expression (dot size) of transcripts of effector/cytotoxic and repair gene signatures for VT samples (n=4). In **B-E**, graphs show combined data for n = 4 for PBMC samples and n = 4 for VT samples.

We predicted that immune cells at mucosal tissue sites must balance host protection with preserving barrier integrity. To understand which cell populations participate in the balance of tissue homeostasis versus inflammatory immune response, we calculated the average expression of curated repair^26^ and cytotoxic/effector gene signatures in our defined cell populations (Figure 1D). Blood effector/memory CD8 T cells and NK cells had the highest effector score of all the populations examined. In contrast, effector/memory CD8 T cells, regulatory T cells (Treg), and NK cell clusters from the VT were each enriched for a tissue repair gene signature (Figure 1D). Since the repair score was high, particularly in immune cells isolated from the vaginal tissue, we analyzed the abundance of individual gene transcripts in immune cells in the VT and found that CD8 T cells and NK cells each score highly on both the effector and repair lists (Figure 1E). To confirm that the repair gene signature was attributed to NK cells, we distinguished the NK cell population from other innate lymphoid cells based on expression of *EOMES* and lack of *CD200R* (Extended Data Figure 1D).^7, 27, 28, 29, 30^ This finding suggests that vaginal tissue NK cells, in addition to an anticipated cytotoxic role in mucosal tissues, may also participate in local tissue repair, thereby playing a dual role in host protection by inducing inflammation and cytotoxicity to reduce pathogen burden and by maintaining tissue homeostasis to protect mucosal tissue integrity.

### Vaginal tissue and circulating NK cells exhibit contrasting functional signatures

The distinct transcriptional signatures of circulating and tissue NK cells (Figure 1C) prompted further characterization of these populations. A differential gene expression analysis on NK cell clusters from PBMC compared to VT revealed 313 genes upregulated in VT NK cells compared to blood NK cells (Supplemental Table 1, Extended Figure 2A). Using published studies defining tissue NK cells ^4, 12, 31^, we compiled an NK cell gene signature and compared blood NK cells to vaginal NK cells (Figure 2A). NK cells from VT compared to the blood were enriched for genes related to tissue residency and immune regulation, while transcripts related to effector and cytotoxic functions were decreased (Figure 2A). Specifically, VT NK cells were enriched in gene transcripts related to tissue residency (*RGS1*, *TCF7*, *SELL, CDCS*), and transcripts related to repair and immune regulation (*TNFAIP3*, *AREG*, *XCL2*). In contrast, many transcripts enriched in blood NK cells were related to cytotoxicity, effector function, and maturation (*GZMB*, *FCGR3A*, *ITGAM*) and were minimally expressed by VT NK cells (Extended Data Figure 2B and Figure 2B).

**Figure 2.**
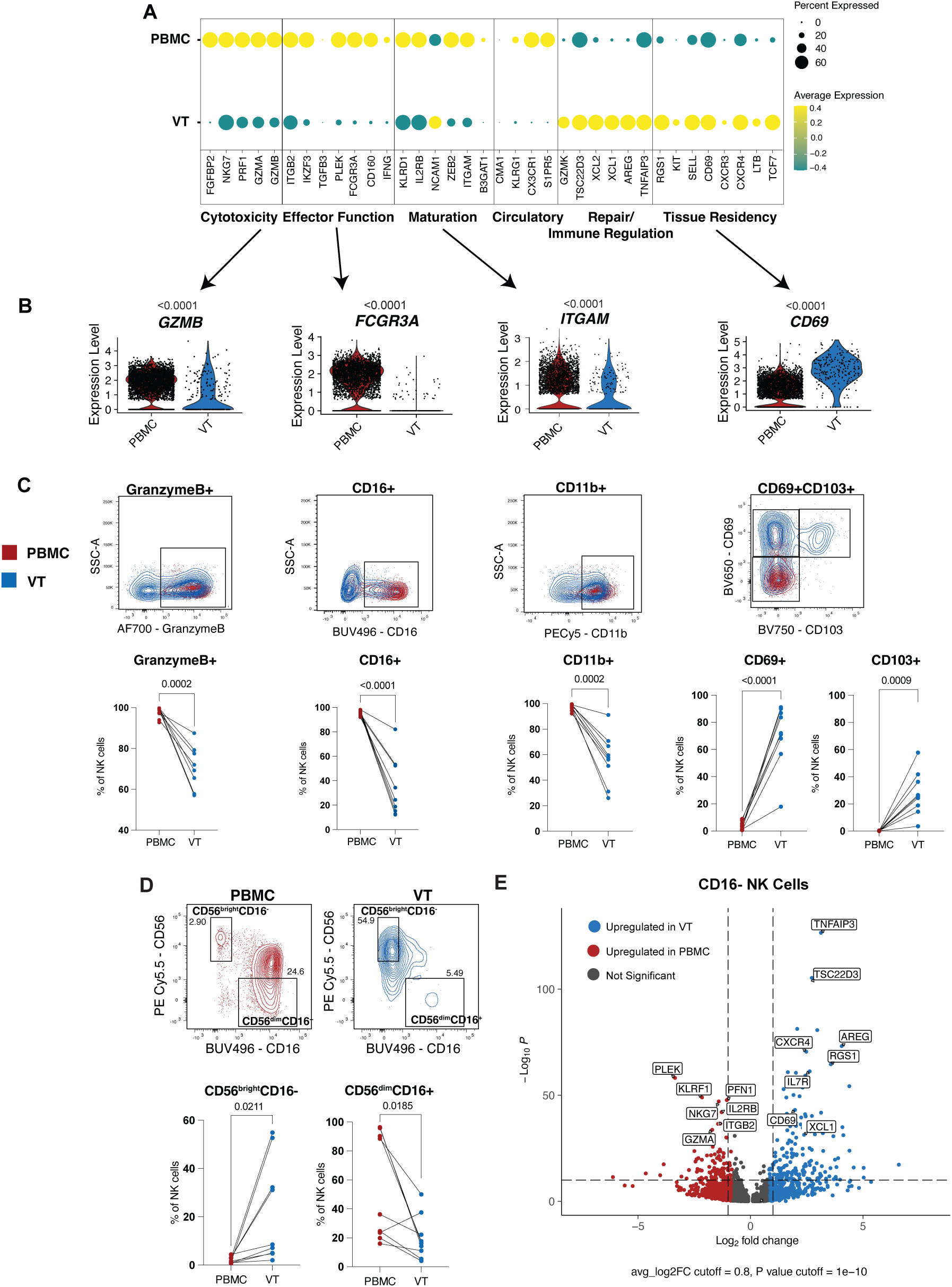
Tissue NK cells and circulating NK cells display differential phenotypic signatures at steady state. Single-cell RNA-seq or flow cytometry was performed on immune cells isolated from paired vaginal tissue (VT) or blood (PBMC) from healthy donors**. A**) Scaled dot plot showing average transcript expression (color) and percent expression (dot size) of select NK cell transcripts from PBMC cluster and VT cluster. **B**) Violin plots showing the expression of selected transcripts for the NK cell clusters from PBMC (left) and the VT (right). All graphs show combined data for n = 4 for PBMC samples and n = 4 for VT samples. **C**) Example flow plot and graphed frequencies of NK cells expressing markers of cytotoxicity, effector function, and tissue residency (combined data for n = 10 for PBMC samples and n = 10 for VT samples). Lines indicate paired samples. **D**) Example flow plots of CD56 and CD16 expression on NK cells from PBMCs (left) or VT (right). Graphs demonstrate frequencies of gated CD56 and CD16 populations. All graphs show combined data for n = 10 for PBMC samples and n = 10 for VT samples. Statistical analyses were performed using paired t test, p values listed. **E**) Volcano plot showing differential gene expression of CD16-NK cell clusters from PBMC vs VT analyzed using the Seurat implementation of MAST (model-based analysis of single cell transcriptomes). Data for n = 4 for PBMC samples and n = 4 for VT samples.

To extend our findings from the transcript to the protein level, we utilized high-parameter flow cytometry to measure markers of NK cell activation, maturation, cytotoxicity, regulation, and tissue residency. Because collagenase digestion has been shown to induce transcriptional changes^32^, this approach also provided orthogonal validation of our transcriptomic data. NK cells were defined as CD45+, CD19-, CD14-, CD3-, and NKp46+, the latter of which was found previously to be a reliable marker for blood and tissue-derived NK cells (Extended Data Figure 2C)^33^. At the protein level, the frequency of NK cells expressing granzyme B, effector molecule CD16, and maturation marker CD11b was decreased in the VT compared to the blood (Figure 2C), consistent with our transcript-level findings (Figure 2B). We measured well-characterized markers of tissue residency, CD69 and CD103 ^4, 5, 7, 9, 34, 35, 36, 37, 38, 39, 40, 41^. These tissue resident markers were also expressed at a significantly higher frequency on NK cells from the VT (Figure 2C). TCF-1, which has been shown to regulate NK cell tissue residency differentiation and be necessary for tissue residency to occur^12^, was also expressed in a greater frequency of VT NK cells (Extended Data Figure 2D). The expression of activating and inhibitory receptors that regulate NK cell function (TIGIT, NKG2A, NKG2D) were increased on NK cells from the VT (Extended Data Figure 2D). We additionally measured markers of activation and maturation (CD57 and CD38) and saw the frequency of NK cells in the VT expressing these markers was significantly decreased when compared to the blood (Extended Data Figure 2D).

Upon examination of markers of NK cell maturation, the frequency of CD56brightCD16-NK cells was higher in the VT, while the frequency of CD56dimCD16+ NK was higher in the blood (Figure 2D). CD56bright NK cells predominate in the lymph nodes and at sites of inflammation, and these NK cells have been shown to have an immune regulatory function^42^ through cytokine production, such as IFNγ, TNFα, and GM-CSF^43^ ^44, 45^. In comparing circulating versus vaginal NK cells, we considered that we could be describing differences between the CD56brightCD16- and CD56dimCD16+ populations since these are differentially enriched in the two compartments (Figure 2D). Using UMAP dimensionality reduction, we clustered the NK cell populations from the PBMC and VT and identified CD16 *(FCGR3A)* negative NK cell populations both in PBMC and VT (Extended Data Figure 2E). We then examined differentially expressed genes in CD16-NK cells and found similar genes related to immune regulation and tissue repair upregulated in CD16-NK cells from the VT compared to CD16-NK cells from the blood (Figure 2E). This analysis suggests that NK cells from the VT are distinct from NK cells from the blood and present a less mature, more regulatory phenotype that may be poised to participate in immunoregulatory functions within tissue sites.

### CD56^bright^ NK cells, found at high frequency in vaginal tissue, produce Amphiregulin

In exploring factors promoting immune homeostasis and tissue protection in the VT, our scRNA-seq data indicated that VT NK cells might be involved in tissue repair functions via expression of Amphiregulin (Areg) (Figure 1E, Extended Data Figure 2B). Areg is a key growth factor that promotes the restoration of tissue integrity following damage associated with inflammation^46, 47, 48^. We confirmed the transcriptional data by measuring Areg expression by NK cells via flow cytometry utilizing a fluorescence minus one (FMO) gating strategy to identify a true positive population (Extended Data Figure 3A). We found a significantly increased frequency of NK cells expressing Areg in tissue NK cells compared to the blood at steady state (Figure 3A). Next, we wanted to determine which subset of NK cells was responsible for Areg production. We found that Areg was predominantly expressed directly *ex vivo* by the CD56^bright^ population of NK cells from the VT (Figure 3B), similar to our finding that *AREG* expression was differentially expressed by vaginal CD16-NK cells compared to circulating CD16-NK cells (Figure 2E).

**Figure 3.**
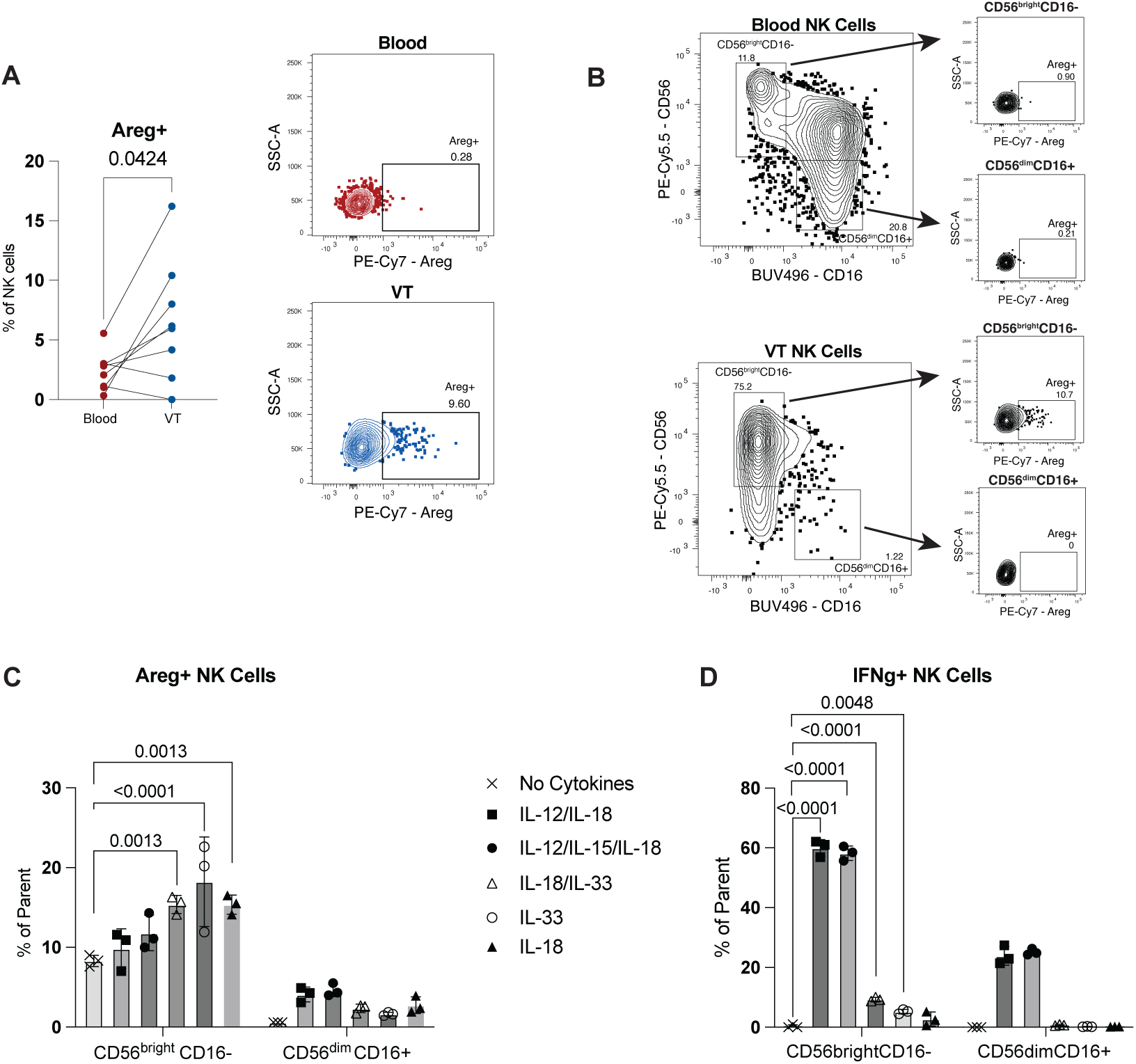
CD56^bright^ NK cells, found in high frequency in vaginal tissue, produce Amphiregulin. Flow cytometry was performed on immune cells isolated from vaginal tissue (VT) or blood (PBMC) from healthy donors. **A**) Paired graph and example flow plot demonstrate frequencies of NK cells expressing Amphiregulin (Areg) in blood (n=8) compared to VT (n=8). **B**) Representative flow plot of CD56^bright^CD16^-^ or CD56^dim^CD16^+^ NK cells expressing Areg. **C-D**) Isolated PBMCs were stimulated with different combinations of cytokines (IL-12 10ng/mL, IL-15 50ng/mL, IL-18 10ng/mL, IL-33 50ng/mL) in complete RPMI media with 50ng/mL of IL-2 for 5 days, and the frequency of NK cells expressing **C**) Areg or **D**) IFNγ by CD56^bright^ and CD56^dim^ populations are graphed. In **A**, statistical analyses were performed using paired t test, p values listed. In **C-D**, data are presented as mean with SD (N=3 donors), and statistical analysis used 2way ANOVA, p values listed.

Next, we investigated factors capable of inducing Areg expression by human NK cells. We hypothesized that NK cells might express Areg under inflammatory conditions to mediate tissue protection in an effort to offset the negative consequences of their cytotoxic function to host tissues. To test this hypothesis, we stimulated NK cells with IL-12, IL-15, and IL-18, cytokines important for activating NK cells downstream of infection^49, 50, 51^, and measured the frequency of Areg expression. We observed that exposure to canonical NK cell activating cytokines (IL-12/IL-15/IL-18) did not significantly increase the frequency of NK cells expressing Areg compared to no cytokines (Figure 3C). We next tested additional combinations of IL-1 family damage-associated alarmins, IL-18 and IL-33. The frequency of NK cells expressing Areg was significantly increased in cells stimulated with IL-33, IL-18, or IL-18/IL-33 together, although the latter did not lead to a synergistic increase in Areg expression compared to either cytokine alone (Figure 3C). We confirmed that Areg expression in circulating NK cells was largely restricted to the CD56^bright^ population (Figure 3C and Extended Data Figure 3B). While combinations of IL-12, IL-15, and IL-18 did not significantly increase Areg expression, they did significantly increase the frequency of CD56^bright^CD16^-^ NK cells producing IFNγ (Figure 3D). Finally, we sought to determine if Areg+ NK cells expressed other effector molecules, and found that NK cells producing Areg were largely a separate population from those expressing IFNγ (Extended Data Figure 3C). This data suggests that different combinations of cytokines found in mucosal tissues can program NK cells to specify their function for pro-inflammatory or tissue repair.

### NK cells from vaginal tissue are poised to respond robustly with anti-pathogen activity upon stimulation

We next wanted to investigate the innate defense function of NK cells in the vaginal mucosa. As this is an important mucosal barrier site, we hypothesized that local immune cells would need to be able to respond quickly to infection or inflammation. However, our transcriptional and flow cytometry data suggested a functionally quiescent profile of NK cells from the healthy VT at steady state. To mimic the induction of inflammation, we stimulated the cells with IL-12 and IL-18 for 24 hours^49, 50, 51^. This stimulation led to a higher frequency of vaginal NK cells expressing the effector molecules granzyme B and IFNγ as well as the upregulation of activation markers CD25 and CD69 (Figure 4A). In addition, we found that a higher frequency of NK cells from the VT expressed IFNγ after stimulation (Figure 4B), and that the VT NK cells expressed higher amounts of IFNγ per cell as measured by median fluorescent intensity compared to blood NK cells (Figure 4C). These data suggest that although the NK cells in the vaginal tissue are relatively quiescent compared to their blood counterparts at steady state (Figure 2A-C), they are able to respond rapidly and robustly to inflammatory stimuli. Finally, markers of tissue residency, immune regulation, and effector function/maturation on NK cells after stimulation, including CD69, ICOS, and CD25, were increased in NK cells from blood and vagina after stimulation (Figure 4D). TCF-1 and NKG2D were decreased in NK cells from the VT after stimulation, while expression in NK cells from the blood remained consistent (Figure 4D). Expression of granzyme B in NK cells from the blood was high at baseline and was not appreciably increased after stimulation (Figure 4D). These data indicate that the functionally quiescent nature of the NK cells from the VT is not due to inherent functional deficiencies, as the NK cells can respond robustly to stimulation, but rather suggests a mechanism of regulation to maintain reduced effector function in the mucosal tissue at homeostasis. We hypothesize that this regulation of tissue NK cells to be effector-like or produce tissue repair factors is crucial to maintaining barrier integrity in mucosal tissue.

**Figure 4.**
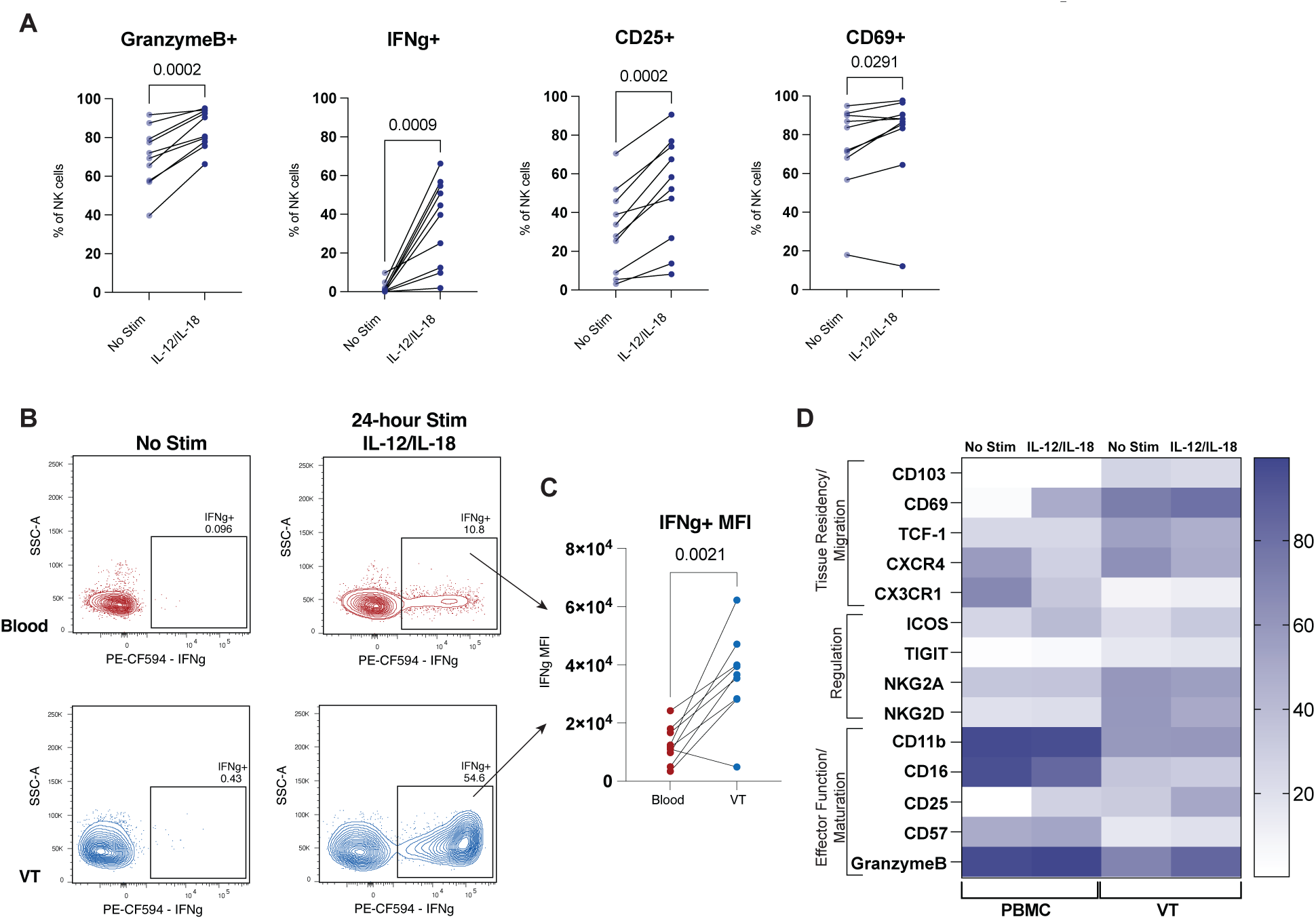
NK cells from vaginal tissue are able to produce an anti-pathogen effector response upon stimulation. Flow cytometry was performed on immune cells isolated from vaginal tissue (VT) or blood (PBMC) from healthy donors and stimulated ex vivo with 10ng IL-12 and 10ng IL-18 or with media alone for 24 hours. **A**) Graphs demonstrate frequencies of NK cells from VT expressing markers of activation and effector function from media alone (n=10) or stimulation with IL-12/IL-18 (n=10). **B**) Example flow plot of NK cell expressing IFNγ under stimulation or no stimulation conditions. **C**) Median fluorescent intensity was calculated from the IFNγ+ population of NK cells after IL-12/IL-18 stimulation from blood (n=9) or VT (n=9) and graphed. **D**) Heat map displaying the mean frequencies of selected markers expressed on NK cells after IL-12/IL-18 stimulation or without stimulation conditions. Statistical analyses were performed using paired t test, p values listed.

### Mouse NK cells recapitulate the human tissue-specific NK cell phenotype

To mechanistically define the factors that regulate the balance of NK cell inflammatory response with tissue repair in a setting of *in vivo* infection, we moved to a well-characterized mouse model of intravaginal herpes simplex virus 2 (HSV-2) infection^52^. To confirm the feasibility of using a mouse model to study NK cells in the VT, we first sorted NK1.1+ NK cells from blood and VT of naïve C57BL/6 mice and performed bulk RNA-seq. Comparing gene expression from VT NK cells and blood NK cells in mice revealed many DEGs (Supplemental Table 2, Extended Data Figure 4A) similar to those found in our human dataset (Supplemental Table 1, Extended Data Figure 2A). In addition, we confirmed our mouse gene expression data by examining protein expression by flow cytometry. To distinguish true tissue-resident NK cells from those merely in circulation, we utilized an intravenous (IV) antibody to label all cells in circulation before VT harvest. This allowed us to analyze NK cells from the VT that were IV label negative and, therefore, resident in the tissue (Extended Data Figure 4B). In mouse NK cells, the markers CD11b and CD27 distinguish stages of NK cell maturation similar to CD16 and CD56 in humans^53, 54, 55^.

CD11b^-^CD27^+^ NK cells are a more immature subset with cytokine secreting potential^54^. This population was significantly more abundant in the VT compared to the blood (Figure 5A). In addition, we saw that CD11b^+^CD27^+^ and CD11b^+^CD27^-^ NK cells, thought to have increased cytotoxic potential,^53, 55, 56^ were significantly higher in the blood compared to the VT (Figure 5A). This led us to hypothesize that similarly to NK cells from the VT in humans, mouse VT NK cells would be immunoregulatory and have increased tissue residency markers.

**Figure 5.**
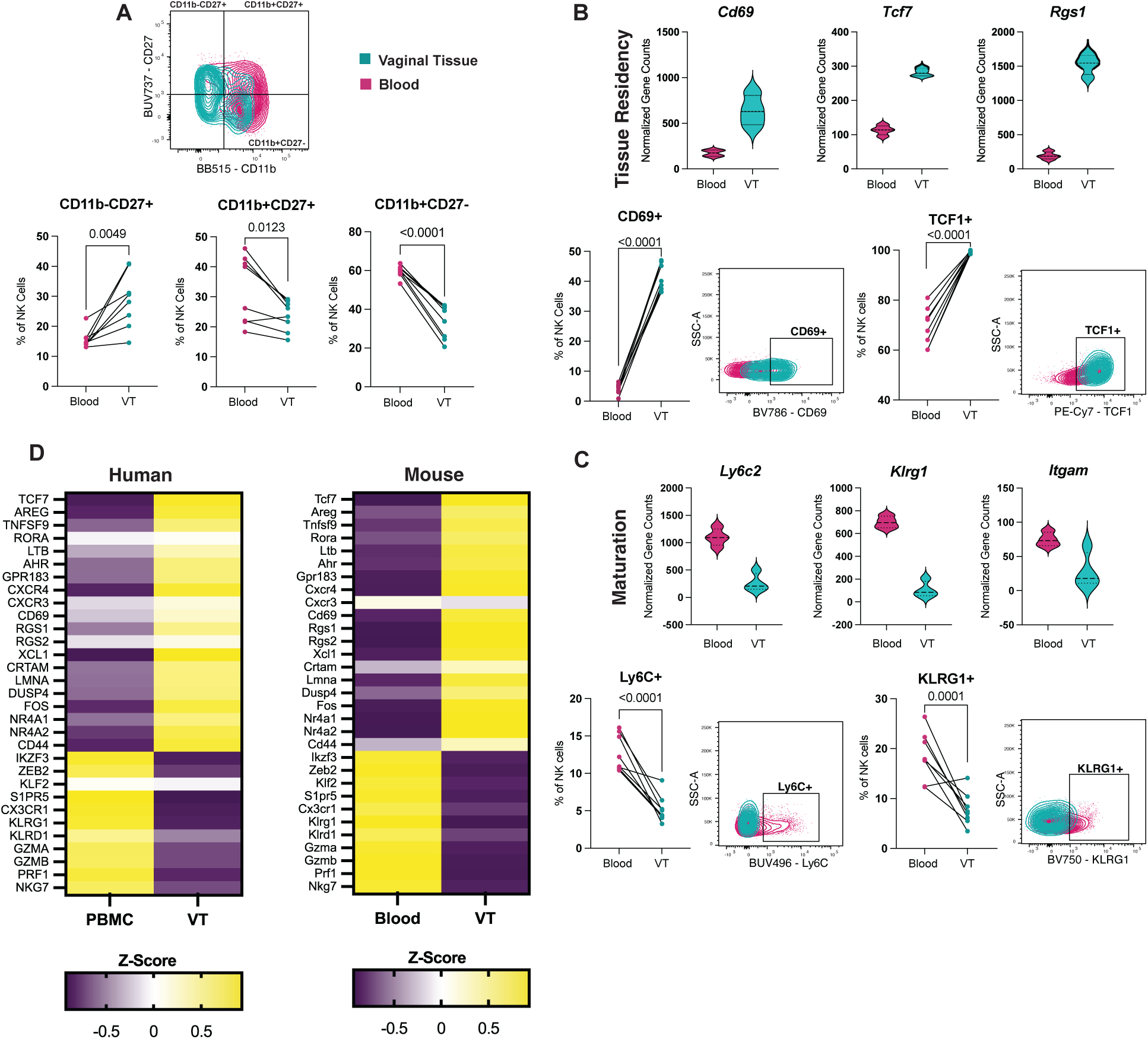
Mouse vaginal tissue NK cells compared to circulating NK cells show similarities to corresponding human NK cell subsets. Vaginal tissues and blood were harvested from uninfected C57BL/6 female mice. NK cells defined as Lymphocytes/SingleCells/Live/ CD45+/Ly6G-/CD19-/CD3-/NK1.1+. Bulk RNA sequencing was performed on sorted NK cells using SmartSeqV4 Ultra Low Input RNA Kit for Sequencing. Flow cytometry was performed on IV Label-NK cells. **A**) Paired graph showing NK cells gated on CD11b and CD27 subsets shown in an example flow plot from blood (n=8) or VT (n=8). **B-C**) Violin plots showing normalized gene counts for selected genes and example flow plots with graphed frequencies from NK cells isolated from blood (left) or VT (right). NK cells expressing markers of **B**) tissue residency and **C**) maturation. RNA-seq data shows n=4 for blood and VT samples. Flow cytometry data shows n=8 for blood samples and VT samples. Flow cytometry data are combined from two independent experiments. **D**) Heat map of calculated z-score for selected genes comparing circulating NK cells to tissue NK cells in both human (left) and mouse (right). Statistical analyses were performed using paired t test, with p values listed.

When we examined the signature of VT NK cells from mice, we found a similar pattern to the human data in that genes responsible for tissue residency, *CdCS*, *Tcf7*, and *Rgs1,* were expressed at higher levels in NK cells from VT compared to the blood (Figure 5B). Concordantly, the frequency of NK cells from the VT expressing CD69 and TCF-1 was significantly higher compared to NK cells from the blood (Figure 5B). Genes related to NK cell maturation and activation *Ly6c2*, *Klrg1*, and *Itgam*^57, 58, 59^ were expressed at lower levels in VT NK cells compared to blood, which was also confirmed by flow cytometry data (Figure 5A,C). We additionally found higher expression of immune regulation and tissue repair genes *Areg*, *Xcl1*, and *Tnfaip3* expressed in NK cells from VT (Extended Data Figure 4C). Genes related to NK cell effector function, *Gzma*, *Gzmb*, and *Prf1,* were expressed at higher levels in blood NK cells compared to VT (Extended Data Figure 4C). Additionally, the frequency of NK cells expressing markers related to tissue retention and immune regulation CD49a, NKG2D, and CD103 were all significantly increased in the VT (Extended Data Figure 4D).

These findings supported those from our human dataset, confirming that both murine and human NK cells from the tissue have a gene signature that suggests less effector function and increased tissue residency and immune regulation at homeostasis. To confirm NK cells from our mouse model were consistent with NK cells from our human samples, we calculated a z-score of gene expression using the cluster average transcript data from human blood and VT NK cells and the bulk RNA-seq data from mouse blood and VT NK cells and compared genes important for NK cell function (Figure 5D). We found that the gene signatures between mouse and human NK cells were remarkably similar, demonstrating the feasibility of using a mouse model of HSV-2 infection to explore vaginal NK cell phenotypes upon local, mucosal tissue infection and inflammation. Overall, this flow cytometry and gene expression data from mouse NK cells mirror human vaginal NK cells in having decreased cytotoxicity and increased tissue retention and immune regulation potential.

### Mucosal NK cells produce cytotoxic and tissue repair molecules during HSV-2 infection

To determine how tissue NK cells alter their function during mucosal tissue infection to balance anti-pathogen activity with tissue repair, we examined NK cells in the VT very early post-infection and at a post-infection timepoint following localized vaginal HSV-2 infection. Mice were synchronized in the diestrous phase of the estrous cycle using Depo-Provera^52^ and then inoculated 7 days later intravaginally with 2×10^6^ PFU of HSV-2 strain 186Kpn TK-^60^ (Figure 6A). The TK^−^ mutant of HSV-2 resolves within 7 days without neurologic disease or death in C57BL/6 mice, allowing us to examine an acute time point with peak viral titers at 2 days post-infection (dpi), and after the viral infection has resolved and during the tissue repair phase, at 10dpi (Figure 6A). To determine how the transcriptional landscape of NK cells is altered throughout the course of infection, we performed bulk RNA-Seq on sorted NK cells from VT at day 0 (uninfected), 2dpi, and 10dpi. Principal component analysis (PCA) of the transcriptional data showed that NK cells from 2dpi clustered separately from NK cells isolated at 10dpi and from uninfected mice based on PC1 (Extended Data Figure 5A). NK cells from uninfected mice and 10dpi clustered separately based on PC2. This suggests that NK cells from uninfected mice and mice 10dpi are more similar than NK cells from mice 2dpi. This was confirmed by differentially expressed gene (DEG) analysis (Extended Data Figure 5B). Specifically, there were 472 significant DEGs (Supplemental Table 3) with an adjusted p-value less than 0.05 comparing NK cells from uninfected VT to 2dpi, indicating a large transcriptional change. Similarly, there were 505 significant DEGs between 2dpi and 10dpi (Supplemental Table 4). However, there were only 11 significant DEGs between uninfected and 10 dpi, suggesting that these populations of cells are more closely related, yet not identical (Extended Data Figure 5B-C).

**Figure 6.**
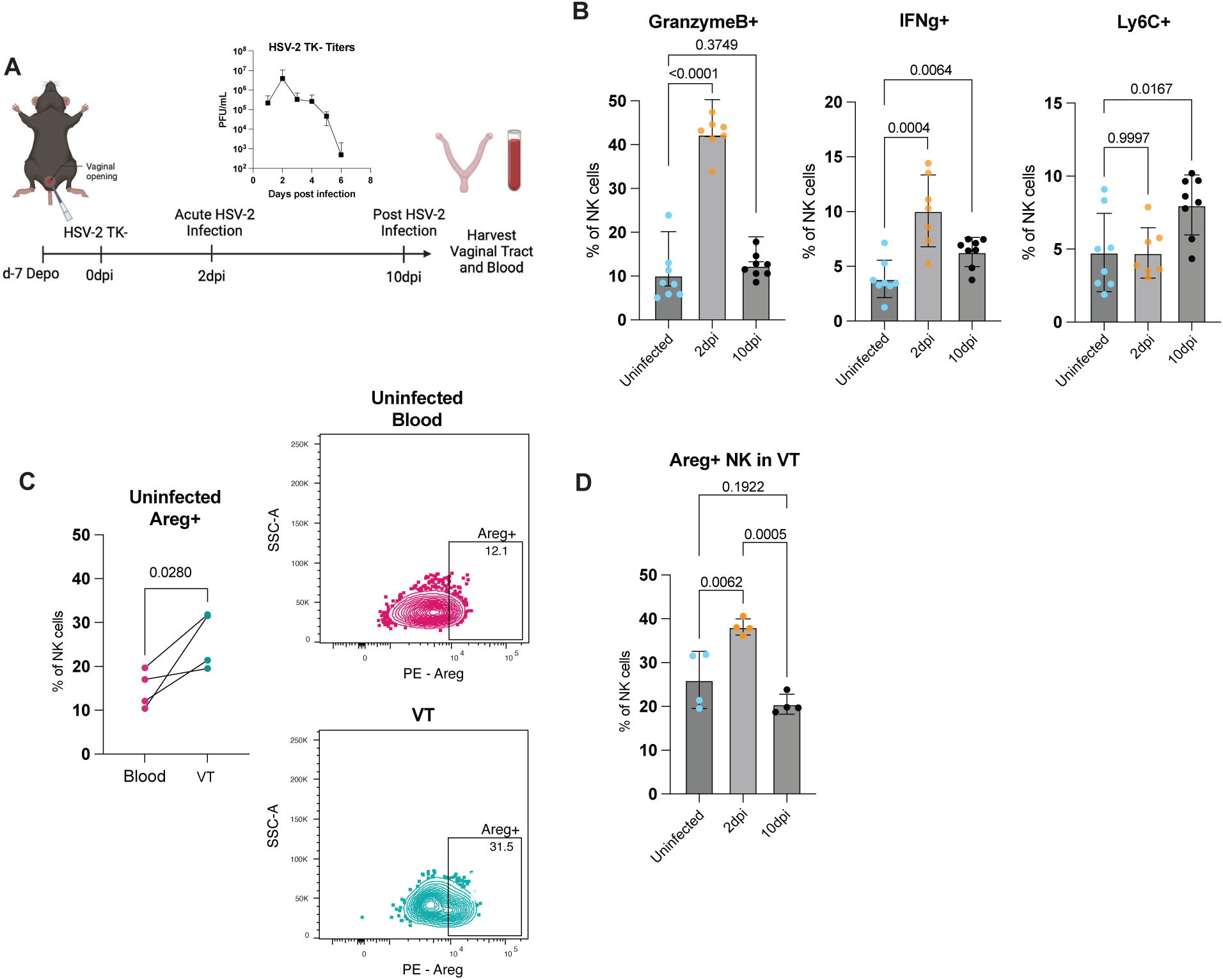
NK cells produce cytotoxic and tissue repair molecules during HSV-2 infection. C57BL/6 female mice were infected vaginally with 2×10^6^ PFU HSV-2 TK- and vaginal tissues and blood were harvested at 2 days post infection (dpi) and 10dpi and from uninfected mice. Flow cytometry was performed on NK cells gated on Lymphoctyes/ SingleCells/Live/ CD45+/Ly6G-/CD19-/CD3-/NK1.1+/IVLabel-. **A**) Experimental design demonstrating strategy for examining NK cells from vaginal tissue or blood at an acute infection time point (2dpi) and post infection time point (10dpi). Viral titers from vaginal washes after HSV-2 TK-infection demonstrate peak infectivity at 2 days post infection and viral clearance by 8 days post infection. **B**) NK cells were stimulated ex vivo with IL-12/IL-18 for 24 hours at 37 °C to measure recall potential. Flow cytometry data indicating frequency of NK cells expressing cytotoxic molecule granzyme B, pro-inflammatory IFNγ, and memory marker Ly6C, from uninfected mice (n=8), 2dpi (n=8) or 10dpi (n=8). **C**) Example flow plot and graphed frequencies of NK cells expressing Areg in uninfected mice comparing blood (n=4) to VT (n=4). **D**) Graphs showing the frequency of NK cells expressing Areg from VT across infection time points (n=4). In **B** and **D**, data are presented as mean with SD, and statistical analysis used multiple unpaired t-test, p values listed. Data are combined from two independent experiments. In **C**, statistical analysis was performed using paired t test, with p values listed. Representative data from two independent experiments.

Specific genes expressed by NK cells at 10 dpi indicated a tissue repair phenotype. Included in the genes unique to NK cells from 10dpi compared to uninfected are *Icos* and *Ly6c2*. Ly6C is thought to be a memory marker on tissue-resident NK cells after viral infection^61, 62^, thereby potentially suggesting that vaginal NK cells retain altered functional capacity following mucosal infection with HSV-2. Additional evidence for induction of an NK cell memory-like population after infection came from upregulation of genes that classify adaptive NK cell populations, including CD3 chain transcripts and members of the histocompatibility gene family, at 10dpi compared to uninfected. We saw genes that define this population reach significance when we reduced the stringency of our adjusted p-value cutoff to 0.1 (Extended Data Figure 5C). To examine the recall potential of NK cells following infection, we collected VT at 2 dpi (acute) and 10 dpi (post-infection) (Figure 6A), stimulated the NK cells *ex vivo* with IL-12 and IL-18 to mimic re-exposure to inflammation, and performed flow cytometry. There was a significant increase in the frequency of NK cells producing granzyme B at 2dpi, with a decrease to homeostatic levels by 10dpi (Figure 6B). IFNγ production followed a similar trend, though IFNγ remained somewhat elevated at 10dpi, suggesting enhanced recall potential of NK cells following infection (Figure 6B). Consistent with this, the frequency of Ly6C+ NK cells was significantly increased at 10dpi compared to uninfected controls, supporting the emergence of a memory-like NK cell population (Figure 6B).

To assess the tissue repair potential of NK cells during HSV-2 infection, we evaluated Areg expression. Under homeostatic conditions, NK cells isolated from the VT express significantly higher levels of Areg than NK cells from the blood (Figure 6C). During infection, Areg expression peaked on NK cells at 2dpi (acute phase) and returned to baseline by 10dpi (post-infection) (Figure 6D). This transient increase in Areg at the height of viral burden suggests that NK cells engage tissue-protective programs to limit immunopathology during pathogen clearance. To directly test this possibility, we assessed the functional impact of NK cell-derived factors on tissue repair.

### NK cell-derived factors directly promote tissue repair

To mimic infection *in vitro* and define the inflammatory cues that induce Areg in murine NK cells, splenocytes were stimulated with combinations of IL-12, IL-15, IL-18, and IL-33, as performed with human cells (Figure 3C). Consistent with the human data, co-stimulation with IL-18 and IL-33 significantly increased the frequency of Areg+ murine NK cells compared to media alone (Figure 7A). To determine whether this effect was NK cell intrinsic, purified splenic NK cells were cultured with IL-18 and IL-33, resulting in detectable Areg production by ELISA (Figure 7B). These findings indicate that NK cells directly respond to inflammatory and alarmin signals present during infection to produce Areg, suggesting a potential role in promoting tissue repair.

**Figure 7.**
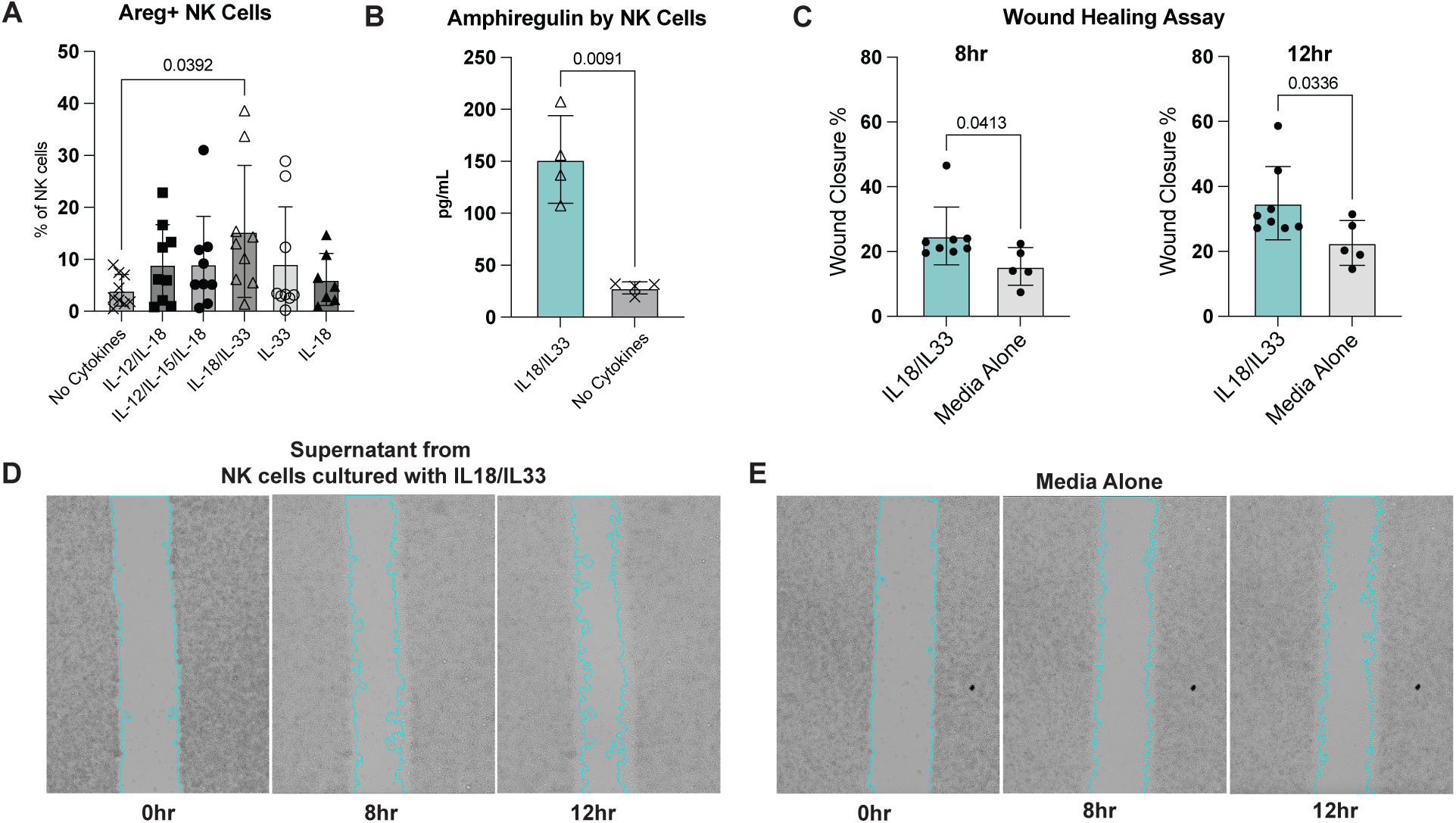
NK cells produce amphiregulin and promote tissue repair. **A**) Splenocytes isolated from naive C57BL/6 female mice were cultured in RP10 supplemented with IL-2 for 24 hours and stimulated with combinations of IL-12, IL-15, IL-18, and IL-33 cytokines. The graph shows the frequency of Areg+ NK cells under each cytokine condition (n=9). Data combined from three independent experiments and presented as mean with SD. Statistical analysis was performed using ordinary one-way ANOVA, p values listed. **B-E)** NK cells were isolated from spleens by negative selection and cultured in DMEM supplemented with IL-2, IL-18, and IL-33 or no cytokines for 40 hours. **B)** Areg levels in culture supernatants were quantified by ELISA. Data are representative of three independent experiments. **C-E)** Conditioned media from NK cell cultures media alone were applied to a mechanically disrupted monolayer of mouse fibroblasts, and wound closure was assessed at 8 and 12 hours. Data are combined from two independent experiments and presented as mean with SD. Statistical analysis was performed using an unpaired t-test, with p values listed.

To test the functional consequence of this response, we assessed whether NK cell-derived factors could directly promote tissue repair using an *in vitro* wound healing assay. Purified NK cells were stimulated with IL-18 and IL-33, and the resulting conditioned media were applied to a mechanically disrupted monolayer of mouse fibroblasts. Wound closure was monitored over time and quantified as the percentage of gap closure. Supernatants from cytokine-stimulated NK cells significantly enhanced wound closure compared to media controls (Figure 7C). Wound area was quantified using an ImageJ-based analysis pipeline ^63^, with images acquired every 4 hours. Enhanced closure was evident as early as 8 hours and persisted at 12 hours in cultures treated with NK cell-conditioned media (Figure 7D), compared to media alone (Figure 7E). The ability of NK cell-conditioned media to promote wound closure *in vitro* suggests that NK cells may contribute to tissue repair during infection, prompting us to assess the impact of NK cell depletion during infection on vaginal tissue barrier integrity.

### Depletion of NK cells disrupts vaginal epithelial barrier integrity and impairs tissue repair during mucosal infection

Finally, we sought to test our hypothesis that NK cells play a direct role in tissue repair in local mucosal tissues during an infection, separate from their role in pathogen clearance. We utilized our intravaginal infection model of HSV-2 and depleted NK cells after the infection had begun to resolve, and then looked at vaginal tissue pathology at 10dpi. Because NK cells are known to be important for the control of HSV-2 infection^64^, we wanted to distinguish the known role of NK cells in viral clearance early post-infection from their potential role in tissue repair later after infection. Based on the viral titers during HSV-2 TK-(Figure 6A), 2dpi is the peak of viral infection, with titers beginning to resolve quickly after 4dpi. Thus, we injected mice with anti-NK1.1 antibody (Ab) to deplete NK cells systemically or IgG2a isotype control Ab on 4dpi, 5dpi, and 8dpi. We then harvested VT at 10dpi for pathology scoring (Figure 8A). NK cell depletion was confirmed in both blood and VT using flow cytometry (Extended Data Figure 6A-B). VT were stained for HCE and scored by a blinded veterinary pathologist in four lesion categories, which included vaginal epithelium, vaginal lamina propria, vaginal muscularis, and vaginal lumen, and received a score of 0-4 for each category (Figure 8B). VT depleted of NK cells after acute HSV-2 infection had significantly increased epithelial damage and lymphocyte infiltration compared to VT from isotype control mice (Figure 8C). The increased pathology occurred despite equal viral clearance: there were no differences in the viral load from vaginal washes between NK cell-depleted and isotype control mice (Extended Figure 6C). To test whether NK cells are a key source of Areg *in vivo*, we quantified Areg+ immune cells (CD45+) in the VT by flow cytometry (Extended Data Figure 6D) at 10 days post-infection following NK cell depletion. NK cell-depleted mice exhibited a marked reduction in Areg+CD45+ cells compared to isotype-treated controls (Figure 8D, Extended Data Figure 6E). Consistent with these cellular findings, Areg levels in vaginal washes, measured by ELISA as an orthogonal approach, were also reduced in NK cell-depleted mice (Figure 8E). Together, these data indicate that NK cells are a critical source of Areg in the VT and likely contribute directly to tissue-reparative processes during HSV-2 infection.

**Figure 8.**
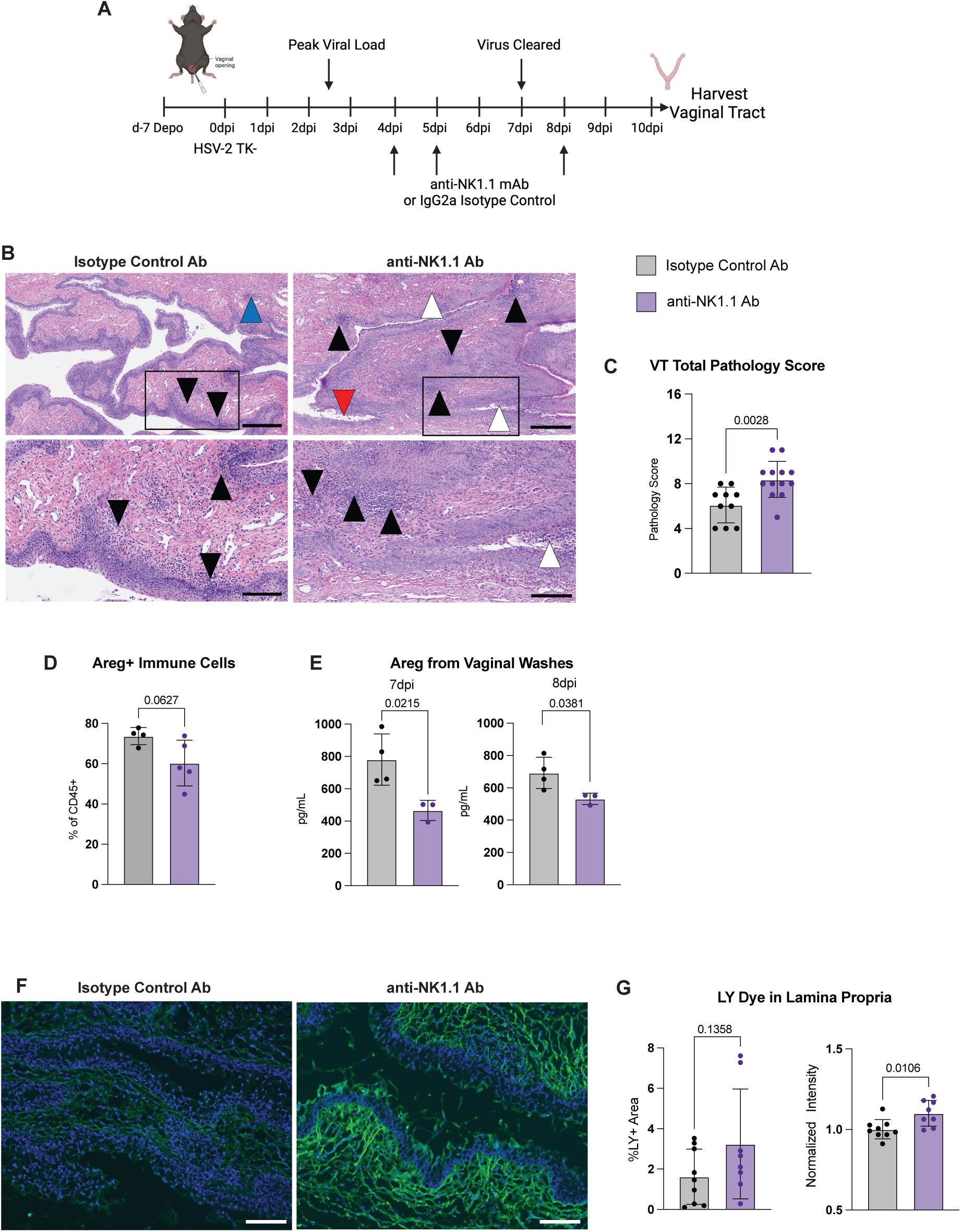
NK cell depletion after HSV-2 infection leads to increased tissue damage. **A**) Experimental model demonstrating mice were infected with 2×10^6^pfu HSV-2 TK- and administered 200ug of NK cell depleting antibody anti-NK1.1 or IgG2a isotype control antibody on days +4, +5, and +8 after infection. Vaginal tissues from mice were harvested on day +10 after infection for HCE staining or flow cytometry. **B**) Representative HCE images from an isotype control (left) and NK cell-depleted (right) vaginal tissue. Arrowheads indicate lesions in the vaginal epithelium (white arrowhead), lamina propria (black arrowhead), muscularis (blue arrowhead) or lumen (red arrowhead). Scale bar 500 uM (top), 100 μM (bottom). **C**) Total pathology scores from HCE images for isotype control (n=10) or NK cell-depleted (n=13) groups. **D)** Frequency of Areg+ of CD45+ cells in mice treated with isotype control (n=4) or anti-NK1.1 (n=5) at 10dpi. Cells were gated on TimeGate/Lymphocytes/SingleCells /CD45+Live /Areg+. Data are combined from two independent experiments. **E**) Areg levels in vaginal washes from NK cell-depleted (n=3) and isotype-treated (n=4) mice were quantified by ELISA at days 7 and 8 post-infection. Data are representative of three independent experiments. **F-G**) Mice were intravaginally administered 50 μg of lucifer yellow (LY) dye 45 minutes before tissue harvest. **F**) Representative fluorescent images of vaginal tissue from isotype-treated (left) and NK cell-depleted (right) mice. Scale bar, 100 μM. **G**) Quantification of the percent of LY+ area and normalized average intensity of LY dye within the lamina propria from isotype-treated (n=9) and NK cell-depleted (n=8) groups. Data are combined from two independent experiments. **C-E, G**) Data are presented as mean with SD, and statistical analysis was performed using an unpaired t-test, p values listed.

Finally, because Areg can act directly to maintain epithelial integrity, we sought to distinguish whether NK cell depletion exacerbates overall inflammation versus directly compromising the vaginal epithelial barrier. To assess barrier integrity, lucifer yellow (LY), a small molecule fluorescent dye, was administered intravaginally shortly before tissue harvest, allowing measurement of dye leakage past the epithelium barrier into the lamina propria^65^. Mice were depleted of NK cells post-acute infection as described above (Figure 8A), and vaginal tracts were harvested 45 minutes after dye administration for imaging. Images were annotated in a blinded fashion by a pathologist, and the area and average intensity of LY in the lamina propria were quantified (Figure 8F). NK cell-depleted mice exhibited greater dye penetration into the lamina propria, both in area and intensity, compared to isotype-treated controls (Figure 8G). Together, our findings reveal an unexpected role for mucosal NK cells in balancing host protection with tissue repair in the VT during localized mucosal infection.

## Discussion

For many infections, the mucosal barrier is the portal of pathogen exposure, yet the regulation of mucosal immunity is poorly understood. NK cells at mucosal sites are well-known to be critical for controlling infections. We show here that they unexpectedly also play a role in protecting this barrier site from tissue damage following mucosal infection. In this study, we profile NK cells from the vaginal tissue of both humans and mice and find a unique phenotype for NK cells at this important barrier site that suggests a unique and previously unexplored functional capacity.

To our knowledge, this is the first study demonstrating a tissue-resident NK cell signature in the vaginal tract. Comparing VT NK cells to tissue-resident NK cells from other sites reveals both shared features and important differences. Dogra et al. profiled human tissue-resident NK cells from non-lymphoid tissues and reported that the lungs were enriched for CD56^dim^CD16^+^ NK cells at levels comparable to blood, with high CD57 expression. In contrast, the VT is enriched for CD56^bright^CD16^-^ NK cells with low CD57 expression. Notably, similar enrichment of CD56^bright^CD16^-^ NK cells has been observed in the gut, where they also display a tissue-resident gene signature, including high CD69 and CD103 expression^4^. These findings suggest that NK phenotypes are shaped by tissue-specific environments and potentially by local functional requirements. In support of this idea, Torcellan et al. found very few tissue-resident NK cells in the skin. However, infection induced an expansion of tissue-resident NK cells, which expressed a gene signature, including Tcf7, closely resembling that of VT NK cells. Finally, NK cells have been well characterized in the uterus owing to their importance during pregnancy. As the VT and uterus are both components of the female genital tract (FGT), we were interested in comparing these two populations. One key difference between NK cells from these distinct FGT anatomical sites is that NK cells from the VT produce IFNγ upon stimulation, whereas NK cells from the uterus have reduced IFNγ production^66^ and have a reduced capacity for cytotoxicity^13, 14^. This important difference most likely represents the different roles these NK cells play in the FGT compartments. While uterine NK cells in the upper FGT most likely play a large role in pregnancy, NK cells in the VT are likely to be more poised to respond to sexually transmitted infection pathogens that represent a larger threat in the lower FGT. Furthermore, maintaining mucosal barrier integrity is likely to be of great importance in the vaginal canal, which faces inflammatory threats including dysbiosis of the vaginal flora, physical and inflammatory trauma from sexual intercourse, and exposure to pathogens. Together, these observations indicate that tissue-resident NK cells adopt both conserved and tissue-adapted programs to balance local immune surveillance and function.

A particularly striking finding from our study was the production of Areg by human and murine NK cells (Figure 3, Figure 6C-D, and Figure 7A-B) and the unexpected role of NK cells in tissue repair (Figure 7C-D and Figure 8). While other immune cells have been shown to play an important role in the repair of tissue after infection, such as ILCs and T cells^67, 68, 69^, this role has not been well elucidated for NK cells. The notion of NK cells as important for tissue integrity is supported by our data, in which NK cell depletion after acute HSV-2 infection leads to increased tissue pathology and barrier disruption (Figure 8). While it is known that NK cells are important during peak HSV-2 infection to manage viral control through production of IFNγ^20, 21, 22^, we have shown that NK cells are also necessary after viral clearance to aid in the restoration of tissue integrity. Previous studies in models of inflammation have suggested that NK cells can limit tissue damage. For example, in a model of corneal epithelial abrasion, NK cells modulate the inflammatory response, and NK cell depletion increased neutrophil infiltration and reduced epithelial proliferation, delaying wound healing^70^. Similarly, in allergic airway inflammation, NK cell depletion impaired clearance of eosinophils and CD4+ T cells, disrupting tissue resolution^71^. These studies highlight an immunoregulatory role for NK cells in controlling inflammation and promoting tissue homeostasis. Our findings extend these observations to the vaginal mucosa during viral infection and reveal that NK cells directly contribute to tissue repair through Areg production. In addition, NK cells can regulate antiviral immune responses by direct elimination of activated CD4 T cells, thereby limiting local inflammation^72, 73^. Consistent with this, NK cell depletion after acute HSV-2 infection led to increased immune cell accumulation in the VT (Figure 8B-C), potentially reflecting unrestrained T cell proliferation. Together, these data establish a critical role for mucosal NK cells in balancing antiviral immunity with tissue protection.

A limitation of the study is that the HSV-2 status of human tissue donors in the study is unknown, and so the study of vaginal tissue NK cells in the context of infection is limited to the study of a mouse model. Because a mouse model of HSV-2 has some key differences from human HSV-2 infection, future studies will involve recruiting women with known HSV-2 serostatus and active viral shedding and lesions to longitudinally evaluate NK cell cytotoxic and tissue protective capacity, as we have done for T cells^74^. Nevertheless, we demonstrate that murine vaginal NK cells share a signature with human vaginal NK cells (Figure 5D), thereby highlighting the applicability of our studies of mouse VT NK cells in the context of human HSV-2 infection.

Additionally, a key area of research moving forward will be to further explore the mechanism whereby NK cells balance the anti-pathogen response with tissue protection and repair. Studies in T cells have demonstrated that Tregs are able to produce Areg after IL-33 and IL-18 stimulation^67^, while memory CD8 T cell populations require TCR stimulation for Areg production^68^. We demonstrate that in NK cells, IL-33 and IL-18 together induce Areg in human and mouse NK cell populations (Figure 3C and Figure 7A), suggesting that there may be unique signals that guide the tissue NK cell response between an anti-pathogen, cytotoxic profile and a tissue healing profile. IL-33 and IL-18 are both members of the IL-1 family and are produced in response to inflammation and HSV-2 infection^75, 76, 77, 78, 79, 80, 81^. A major unanswered question is whether there is a tissue-specific signal allowing for this balance of function. Future studies will explore the dynamics of cytokine signaling during infection that promotes the dual nature of NK cell function to transition from protection from pathogens to tissue protection.

In sum, we hereby characterize circulating and vaginal tissue NK cells from human and mouse and demonstrate a transcriptional and protein signature for NK cells from the lower female genital tract. These tissue NK cells are maintained in a functionally quiescent profile at steady state, yet are poised to respond robustly with anti-pathogen activity to inflammatory signals from tissue infection. We additionally show that mucosal NK cells can secrete the tissue repair factor Amphiregulin and play a critical tissue protective function after resolution of infection. We propose a novel mechanism whereby tissue NK cells mediate an active balance between anti-pathogen effector function and preservation of host tissue integrity. Thus, mucosal NK cells are a double-edged sword in the context of viral infection and provide a balance between pathogen clearance and barrier homeostasis.

## Methods

### Human Tissue Collection and Processing

Human blood and VT were collected from women undergoing vaginal reconstructive surgeries for pelvic organ prolapse. Any samples suspected of malignancy were excluded from collection. All participants signed a written informed consent prior to inclusion in the study, and the protocols were approved by the institutional review board at the Fred Hutchinson Cancer Center and the University of Washington (IRB 4323). Blood was collected into sodium heparin tubes (BD Vacutainer) and mixed. Blood was diluted 1:2 in PBS, layered on a Lymphoprep (StemCell Technologies) gradient and centrifuged at 1200 rpm for 20 min. The PBMC layer was then removed and washed with PBS. Tissues were transported from the clinic in ice cold PBS and immediately processed upon arrival (within 1–2 h of removal). Vaginal tissues were trimmed to 2 mm of epithelium and the outermost stroma. Tissue was minced and pipetted into a 50mL Falcon tube with 30 mL of digestion medium and shaken for 30. Minutes at 37 °C. Digestion medium consisted of collagenase II (700 units/ml, Sigma-Aldrich) and DNAse I (1 unit/ml, Sigma-Aldrich) in a 1:1 mixture of PBS and R15 as previously described^82^. The sample was then aspirated and expelled from a 30 mL syringe through a 16-gauge blunt needle and strained through a 70-um strainer. This process was repeated with the remaining tissue pieces and then cells were pelleted and resuspended in fresh RP10 medium.

### Human Cell Culture

Directly following isolation or thawing, cells were cultured with recombinant human IL-12 (Preprotech) at 10ng/mL and recombinant human IL-18 (Peprotech) at 10ng/mL for 24 hours in RP10 at 37°C and 5% CO2. Brefeldin A was added in the last 4 hours of culture. For some experiments, cells were cultured in IL-12 (10ng/mL), IL-18 (10ng/mL), IL-15 (50ng/mL, Peprotech), or IL-33 (50ng/mL, RCD Systems) in RP10 containing 50ng/mL IL-2 (Peprotech) and cultured for 5 days at 37°C and 5% CO2.

### Flow Cytometry

For flow cytometric analysis good practices were followed as outlined in the guidelines for use of flow cytometry^83^ and consensus suggestions for data analysis^84^. Directly following isolation or thawing, cells were incubated with Fc-blocking reagent (BioLegend Trustain FcX) and fixable UV Blue Live/Dead reagent (ThermoFisher) in PBS (Gibco) for 20 min at room temperature. After this, cells were incubated for 20 min at room temperature with 50 μl total volume of antibody master mix freshly prepared in Brilliant staining buffer (BD Biosciences), followed by two washes in fluorescence-activated cell sorting (FACS) buffer (PBS with 0.5% FBS). All antibodies were titrated and used at optimal dilution, and staining procedures were performed in 96-well round-bottom plates. The stained cells were fixed with eBioscience Foxp3 Transcription Factor Fixation/Permeabilization Kit (Thermo Fisher) for 30 min at room temperature, and then stained for intracellular (IFNg, granzyme B (GzmB), or Areg) or intranuclear staining (Ki67 or TCF1) in 50 μl total volume of antibody master mix for 30 min. Samples were stored at 4°C in the dark until run on a flow cytometer. Single-stained controls were prepared with every experiment using antibody capture beads (OneComp eBeads, Thermo Fisher) diluted in FACS buffer, or ArC Amine Reactive Compensation beads (Thermo Fisher) for Live/Dead reagent. All single stain controls were treated the same as the samples (including fixation procedures).

All samples were acquired using a FACSymphony A5 (BD Biosciences) and FACSDiva acquisition software (BD Biosciences). Detector voltages were optimized using a modified voltage titration approach^85^ and standardized from day to day using MFI target values and 6-peak Ultra Rainbow Beads^86^ (Spherotec). After acquisition, data was exported in FCS 3.1 format and analyzed using FlowJo (version 10.6.x, and 10.7.x, BD Biosciences). Samples were analyzed using manual gating with doublets being excluded by FSC-A vs FSC-H gating. Gates were kept the same across all samples except where changes in the density distribution of populations clearly indicated the need for sample-specific adjustment.

### Single-cell RNA-Seq

Single-cell RNA-seq was performed on sort-purified CD45+ and NKp46+ NK cells derived from either cryopreserved blood or VT tissue samples. In total, 8 samples were sequenced from 4 donors.

### Whole transcriptome single-cell library preparation and sequencing

Cryovials of PBMC and VT cells were thawed and added to prewarmed RP10 media and washed. VT cells were enriched for CD45+ cells (EasySep Release Human CD45 Positive Selection Kit, Stemcell) and all samples were stained with Live/Dead and FcBlock for 20 min at room temperature. Cells were resuspended in 80µl of antibody cocktail and 20µl Sample Tag and incubated for 20 min at room temperature. Cells were washed and resuspended in RP10. Cell sorting was performed on a FACSymphony S6 cell sorter (BD Biosciences) with a 70-μm nozzle at 70 psi sheath pressure into Eppendorf tubes with 300 µl RP10. Total CD45+ cells were sorted until around 90% of the desired number cell number was reached, and then NKp46+ cells were sorted until a total desired cell number was reached, or until sample end.

After sorting, cells were pooled, counted, and loaded onto a nano-well cartridge (BD Rhapsody), lysed inside the wells followed by mRNA capture on cell capture beads. Cell Capture Beads were retrieved and washed prior to performing reverse transcription and treatment with Exonuclease I. Libraries were prepared using mRNA whole transcriptome analysis and sample tag library preparation (BD Rhaspsody)^87^. Briefly, mRNA and Sample-Tag were amplified separately and purified using double-sided size selection using SPRIselect magnetic beads (Beckman Coulter). Quality of PCR products was determined by using an Agilent 2200 TapeStation with High Sensitivity D5000 ScreenTape (Agilent) in the Fred Hutch Genomics Shared Resource laboratory. The quantity of PCR products was determined by Qubit with Qubit 1x dsDNA HS Assay (Q33231). Final libraries were diluted to 4nM and multiplexed for paired-end sequencing on a NovaSeq 6000 (Illumina) using an S1 flow cell. For the mRNA library, we targeted 30,000 reads per cell, and for the Sample-Tag libraries, 1,200 reads per cell.

### Pre-processing for whole transcriptome analysis (WTA) and Seurat workffow

Fastq files were processed via the standard Rhapsody analysis pipeline (BD Biosciences) on Seven Bridges (www.sevenbridges.com). In brief, after read filtering, reads are aligned to a reference genome and annotated, barcodes and UMIs are counted, followed by determining putative cells. The final output was analyzed in R using the package Seurat^88^ (version 4.0), with scripts based on the following general guidelines for analysis of scRNA-seq data^89^. In brief, for whole-transcriptome data, only cells that had a valid Sample Tag, at least 200 genes and less than 35% mitochondrial genes were included in analysis. All acquired samples were merged into a single Seurat object, followed by a natural log normalization using a scale factor of 10,000, determination of variable genes using the vst method, and a z-score scaling. Principal component analysis was used to generate 75 principal components and the first 25 were used to calculate UMAP, and graph-based clustering with a resolution of 0.5. For cell annotation, we applied SingleR as a purely data-driven approach^90^, and used the expression of typical lineage transcripts to verify the cell label annotation.

For all differential gene expression analyses we utilized the Seurat implementation of MAST (model-based analysis of single-cell transcriptomes). For calculating the Repair scores we used the AddModuleScore function of Seurat. The genes used were as follows: Effector: GZMB, GZMA, PRF1, NKG7, FGFBP2, GNLY, TNF, IFNG; Repair: IL10RA, HIF1A, CD44, LYST, TNFAIP3, AREG, TSC22D3, TGFB1.

### Mice

WT mice on a C57BL/6 background were used. All animal experiments were approved by the IACUC review board at the Fred Hutchinson Cancer Center (D16-001-42) The office of Laboratory Animal Welfare of NIH approved Fred Hutch (#A3226-01), and this study was carried out in strict compliance with the PHS policy on Humane Care and Use of Laboratory Animals.

### HSV-2 Infection and Viral Titers

HSV-2 186DeltaKpn TK-strain (HSV-2 TK)^91^ was grown and titered on Vero cells. Mice were subcutaneously injected with 2 mg of medroxyprogesterone acetate (MDPA) injectable suspension (Depo-Provera) dissolved in sterile PBS 5–7 days before vaginal infection. For HSV-2 infection, mice were infected intravaginally with 10^6^ PFU of HSV-2 TK-. To determine viral titers, vaginal washes were performed daily after infection with 50 μL titration buffer (PBS + 1% FBS + 1% Glucose + 0.01% MgCl + 0.01% CaCl). One mL of diluted vaginal washes were added to confluent flasks of Vero cells for 1 hour at 37°C, then human IgG at 4.8mg/ml was added to prevent further infection. Vero cells were incubated at 37°C and 5% CO2 for 48 hours and then plaques were counted to determine plaque forming units per mL.

### Mouse Tissue Collection and Processing

The vaginal tissue and cervix were harvested and minced thoroughly in prewarmed digestion medium, which was freshly prepared and consisted of DMEM, collagenase D (2 mg/mL; Sigma-Aldrich), 500 μL DNase (15 μg/mL), and 10% FBS. The minced tissue was incubated for 30 minutes at 37°C on a shaker. After incubation, the minced tissue was spun at 1200 rpm for 5 minutes. To prepare single-cell suspensions, 5 mL of HBSS/EDTA solution (2% FBS in HBSS without calcium and magnesium and 5 mM EDTA) was added to the pellet, and the mixture was homogenized through a 70 μm strainer. Blood was collected through cardiac puncture after euthanasia into Heparin in PBS. ACK lysis buffer was added to the blood for 15 minutes and centrifuged at 300 *g* for 5 minutes. The ACK-treated cells were suspended in HBSS/EDTA solution before proceeding with flow staining.

### Mouse Cell Culture and Cell Depletion

Immune cells isolated from the VT or blood were cultured in round-bottom 96-well plates with 20ng/mL recombinant mouse IL-12 (BioLegend) and 10 ng/mL recombinant mouse IL-18 (BioLegend) in RP10 with BrefeldinA at 1:1000 dilution for 4 hours at 37°C at 5% CO2. For some experiments, splenocytes were cultured in IL-12 (20ng/mL), IL-18 (10ng/mL), IL-15 (20ng/mL, Peprotech), or IL-33 (50ng/mL, RCD Systems) in RP10 containing 20ng/mL IL-2 (Peprotech) and cultured for 24 hours at 37°C and 5% CO2. NK cells were depleted using InVivo anti-mouse NK1.1 clone PK136 (BioXCell). Anti-NK1.1 mAb or IgG2a Isotype Control mAb (BioXCell) were injected intraperitoneally at a concentration of 200ug in 200ul on days 4, 5, and 8 after HSV-2 TK-infection.

### Ampheregulin Quantification by ELISA

Amphiregulin was detected using the Mouse Amphiregulin DuoSet ELISA (RCD Systems) following the manufacturer’s protocol. Conditioned media from isolated NK cells cultured in IL-18 (10ng/mL), IL-33 (50ng/mL, RCD Systems) and 20ng/mL IL-2 (Peprotech) in DMEM with 1% FBS were used to measure Areg produced directly by NK cells. DMEM +1% FBS was used as a background control. To determine Areg production in vivo after NK cell depletion, vaginal washes were taken from NK cell-depleted or isotype-treated mice on days 7 and 8 post-infection, washing with 50 μL of PBS into the vaginal tract of mice before diluting 1:5 in PBS.

### Bulk RNA-seq

Bulk RNA-seq was performed on 250 sort-purified NK1.1+ NK cells derived from either blood or VT tissue samples harvested from uninfected mice or from mice infected with HSV-2 TK-on days 2 and 10 post-infection. In total, 24 samples were sequenced, and each condition was represented by 4 biological replicates.

Cell sorting was performed on a Sony MA900 (Sony) using a 100-μm sorting chip with 90% sort efficiency and 90-95% purity. Cells were sorted directly into reaction buffer from the SMART-Seq v4 Ultra Low Input RNA Kit for Sequencing (Takara), and reverse transcription was performed followed by PCR amplification to generate full-length amplified cDNA. Bulk RNA-seq was performed by the Genomics Core at Benaroya Research Institute.

Sequencing libraries were constructed using the NexteraXT DNA sample preparation kit with unique dual indexes (Illumina) to generate Illumina-compatible barcoded libraries. Libraries were pooled and quantified using a Qubit® Fluorometer (Life Technologies).

Sequencing of pooled libraries was carried out on a NextSeq 2000 sequencer (Illumina) with paired-end 59-base reads (Illumina) with a target depth of 5 million reads per sample. Base calls were processed to FASTQs on BaseSpace (Illumina), and a base call quality-trimming step was applied to remove low-confidence base calls from the ends of reads. The FASTQs were aligned to the GRCm38 mouse reference genome, using STAR v.2.4.2a and gene counts were generated using htseq-count. QC and metrics analysis was performed using the Picard family of tools (v1.134). Filtered and normalized gene counts were generated from the raw counts by trimmed-mean of M values (TMM) normalization and filtered for genes that are expressed with at least 1 count per million total reads in at least 10% of the total number of libraries. To detect differentially expressed genes between sorted cell subsets, the RNA-seq analysis functionality of the linear models for microarray data (Limma) R package was used^92, 93, 94^. Expression counts were normalized using the TMM algorithm^95^. A false discovery rate adjustment was applied to correct for multiple testing.

### Wound Healing Assay

NK cells were isolated from the spleens of naïve females on a C57BL/6 background (8-12 weeks) using an NK cell isolation Kit (Miltenyi Biotec) following the manufacturer’s protocol. Purified NK cells were then cultured in DMEM supplemented with 1% FBS and IL-18 (10ng/mL), IL-33 (50ng/mL, RCD Systems), and 20ng/mL IL-2 (Peprotech) for 40 hours at 37°C and 5% CO2. Conditioned media were collected and spun down to remove cellular debris. Mouse fibroblast L929 (ATCC) cells were seeded in a 12-well plate at 1×10^5^ cells/mL and cultured overnight until 80% confluence. Cell monolayers were scratched with a 200 μL sterile pipette tip. Wells were then washed with PBS (1X) before 1mL of conditioned media from the NK cell culture or DMEM supplemented with 1% FBS (media alone) was added. Brightfield images of wells were taken using a Keyence BZ-X800 microscope at X4 magnification. Fixed points were set for each well, and automated multi-point acquisition was enabled to capture time-lapse images at 0, 4, 8, 12, and 24 hours. Images were analyzed using *Wound Healing Size Tool^C3^*, an ImageJ/Fiji plugin that allows the quantification of wound area, wound coverage of total area, average wound width, and width standard deviation. Percent wound closure was calculated at each time point by subtracting the open area measured in inches^2^ from the open area at the 0-hour time point, divided by the area at the 0-hour time point, multiplied by 100.

### H&E & Whole Slide Digital Imaging

Hematoxylin and Eosin (HCE) staining and whole slide digital imaging was performed by the Fred Hutchinson Cancer Center Experimental Histopathology Shared Resource (P30 CA015704). Tissues were formalin-fixed and paraffin embedded on an automated tissue processor (Sakura Tissue-Tek VIP), sectioned at 4-5 um onto charged slides, baked at 60°C, stained using an automated stainer (Sakura Tissue-Tek Prisma) for HCE staining and cover slipped with permanent mounting media. Whole slide digital imaging was performed on an Evident VS200 at 20X.

### Histopathology Scoring

Slides were examined by a board-certified veterinary pathologist. Digital slides were annotated to exclude cervical, uroepithelial and uterine tissue from grading. Each vaginal tract was examined in four lesion categories which included vaginal epithelium, vaginal lamina propria, vaginal muscularis, and vaginal lumen, and received a score of 0-4 for each category. *Vaginal Epithelium* Score 0: No lesions, Score 1: Epithelial surface is multifocally tattered, rare leukocytes and/or neutrophils within the epithelium, Score 2: Clusters of neutrophils within the epithelium, Score 3: Epithelium is thin and eroded, Score 4: Epithelium is ulcerated; *Vaginal Lamina Propria and Vaginal Muscularis* Score 0: No lesions, Score 1: Rare neutrophils and/or leukocytes, Score 2: Small clusters of neutrophils and/or leukocytes, Score 3: Multifocal clusters of neutrophils and/or leukocytes beginning to coalesce and/or forming follicles, Score 4: leukocytes arranged in sheets; *Vaginal Lumen* Score 0: No lesions, Score 1: Rare cellular debris in lumen, Score 2: <10% of vaginal lumen contains cellular debris, Score 3: 10-25% of vaginal lumen contains cellular debris, Score 4: >25% of vaginal lumen contains cellular debris.

### Vaginal Tissue Permeability Assay

Female BALB/c (8-12 weeks old) were intravaginally inoculated with 50 ug of lucifer yellow CH, <u>lithium salt</u> (MW = 457.2Da) (Cat# L453, ThermoFisher) in PBS. After 45 minutes, mice were euthanized and vaginal tracts were harvested, and placed in 4% paraformaldehyde (Cat #3190-32 Thermo Fisher Scientific) for 4-8 hours, washed in PBS, embedded in OCT compound (Thermo Fisher Scientific), and stored at 80°C. Frozen tissues were sectioned using a cryostat at a thickness of 5 um and mounted onto glass slides. Endogenous peroxidase was blocked with 3% H2O2 for 5 minutes followed by protein blocking with TCT buffer (0.05M Tris, 0.15M NaCl, 0.25% Casein, 0.1% Tween 20, 0.05% ProClin300 pH 7.6) for 10 minutes. Slides stained with DAPI (Sigma-Aldrich D8417-10mg) for 5 minutes, rinsed for 5 minutes, and coverslipped with Prolong Gold Antifade reagent (Invitrogen/Life Technologies, Grand Island, NY). Whole slide images were acquired on the Evident VS200 at a 20x magnification. HALO (Indica Labs, Albuquerque, NM) was used for digital image analysis. Digital images of the vaginal tract were annotated by a board-certified pathologist. The HALO Area Quantification FLv3.0.1 algorithm was used to quantify infiltration of the lucifer yellow (LY) dye in the lamina propria based on the fluorescent intensity of the LY dye in the FITC channel. A mean intensity threshold above background was used to determine positivity of the dye, thereby defining an area as either positive or negative. The positive staining data was then used to quantify the area of LY dye per pathologist-annotated region. To account for inter-experimental variability in fluorescence intensity measurements, values were normalized within each experiment to the mean WT control.

## Acknowledgments

We thank all members of the Lund and Prlic labs for their helpful discussions on experimental findings and manuscript preparation throughout this process. We thank Experimental Histopathology Shared Resources for technical assistance and support. This work was supported by the following grants from the National Institutes of Health: R01 AI131914 (to JML), R01 AI141435 (to JML), R01 AI172111 (to FH and JML) and R01 HD114505 (to JML). SCV was supported by T32 AI007509 and K99 AI180649-02. LSW was supported by T32 AI083203. This research was supported by the Experimental Histopathology Shared Resource, RRID:SCR_022612 and Comparative Pathology Shared Resource RRID:SCR_022610 of the Fred Hutch/University of Washington/Seattle Children’s Cancer Consortium (P30 CA015704).

