## Extended Data Figures for "Mucosal tissue NK cells directly mediate tissue protection and repair during infection"

**Extended Data Figure 1.**

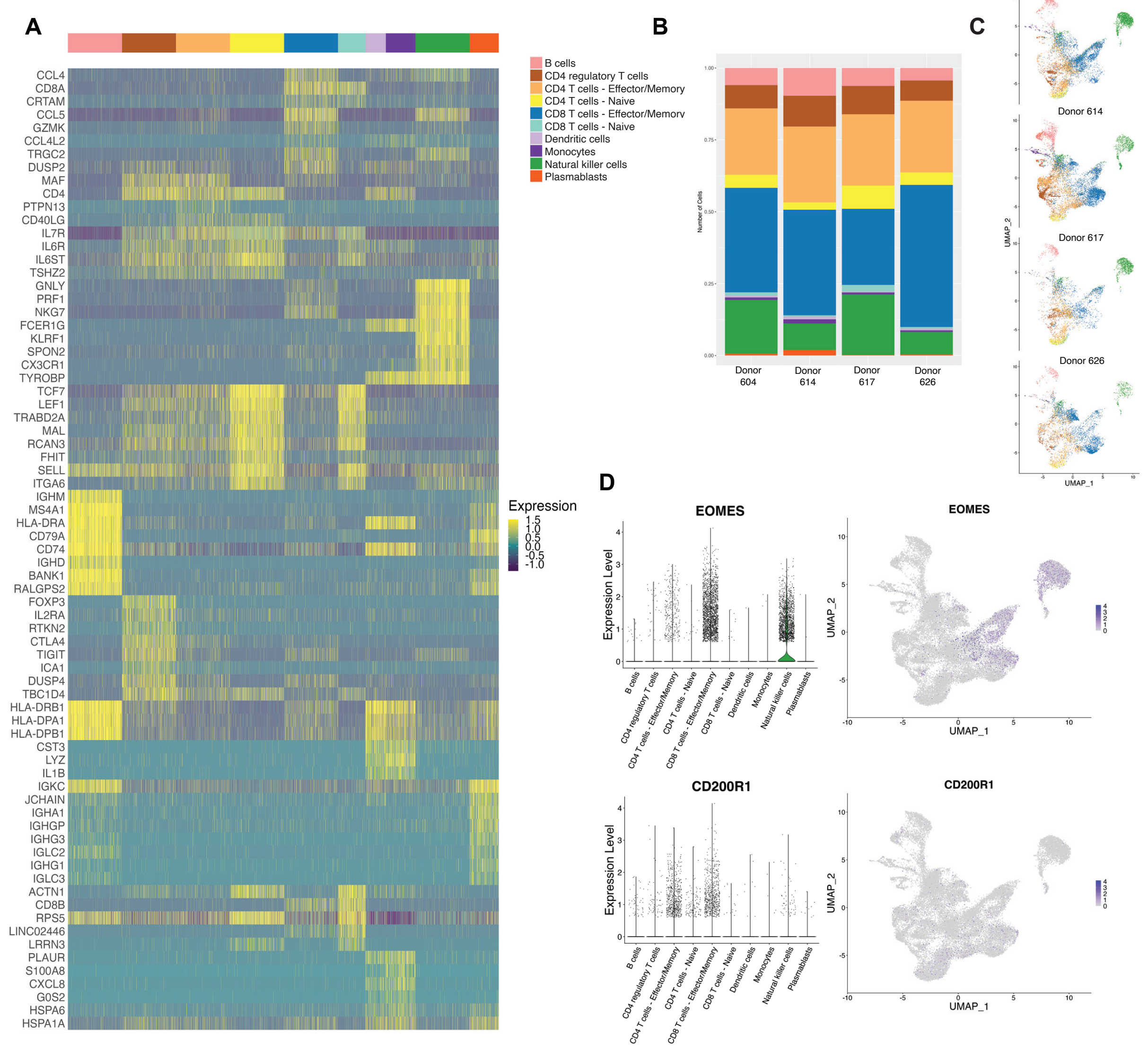


**Extended Data Figure 1. A**) Heatmap showing the top 8 differentially expressed genes across all clusters. **B**) Stacked bar graph showing the makeup of immune cell populations by donor. **C**) UMAP plot of clusters separated by donor. **D**) Violin plot of transcript expression for EOMES and CD200R with expression overlaid on UMAP of immune populations. Data shows combined PBMC and VT data for all donors.

### Extended Data Figure 2.

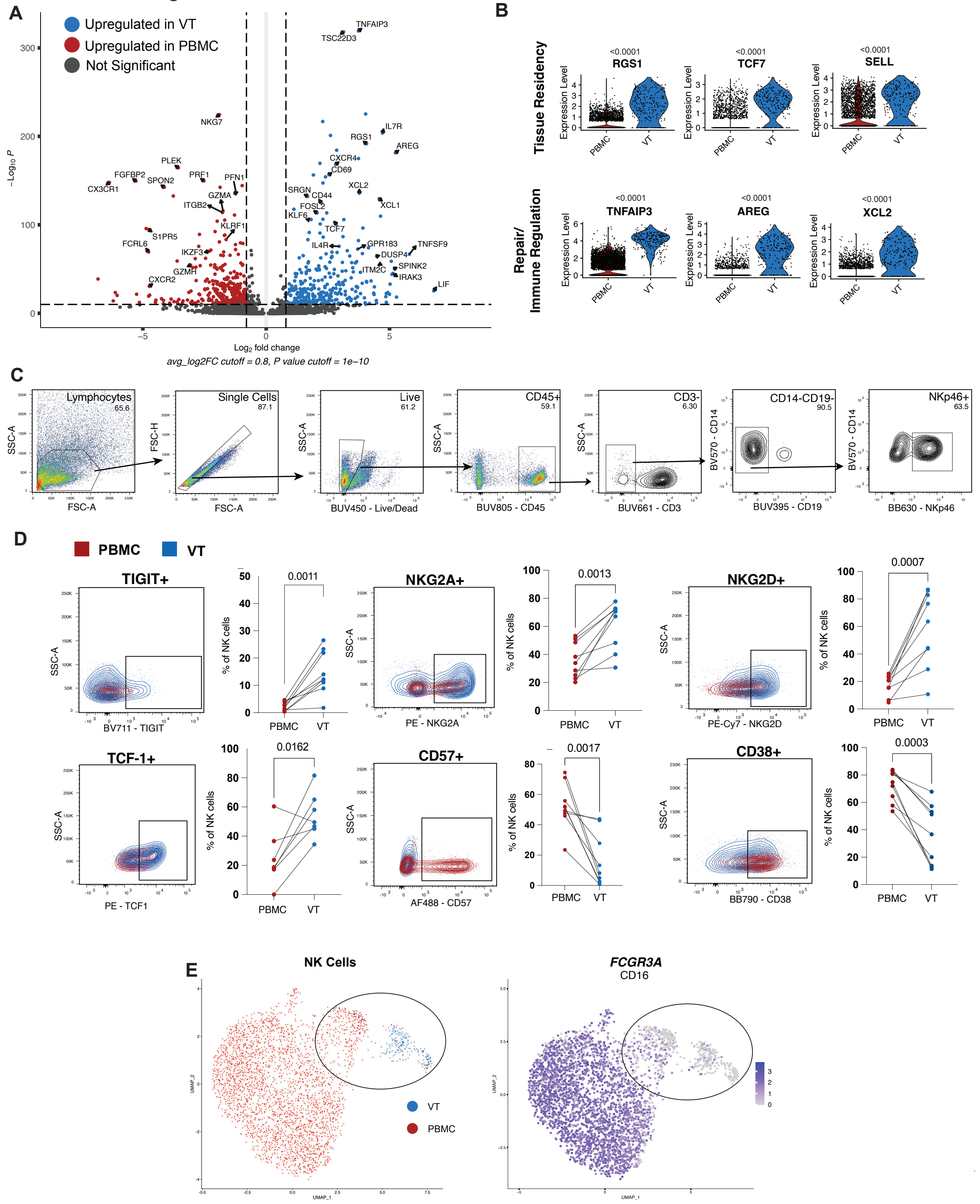


**Extended Data Figure 2. A**). Volcano plot showing differential gene expression of NK cell clusters from PBMC vs VT analyzed using the Seurat implementation of MAST (modelbased analysis of single cell transcriptomes). **B**) Violin plots showing the expression of selected transcripts related to tissue residency and immune regulation for the NK cell clusters from PBMC (left) and the VT (right). **C**) Flow cytometry was performed on immune cells isolated from paired vaginal tissue (VT) or blood (PBMC) from healthy donors**.** Cells were stained for flow cytometry, and data was collected on FACS Symphony. NK cells were gated on TimeGate/Lymphocytes/SingleCells/ Live/CD45+/CD3-/CD19-CD14-/NKp46+. **D**) Example flow plot and graphed frequencies of NK cells expressing markers of regulation and activation/maturation. All graphs show combined data for n = 10 for PBMC samples and n = 10 for VT samples. Statistical analyses were performed using paired t test, p values listed. **E**) UMAP plot based on PBMC and VT NK cell clusters overlaid with FCGR3A (CD16) expression. The black circle indicates populations used to calculate differentially expressed genes in F. All graphs show combined data for n = 4 for PBMC samples and n = 4 for VT samples.

**Extended Data Figure 3.**

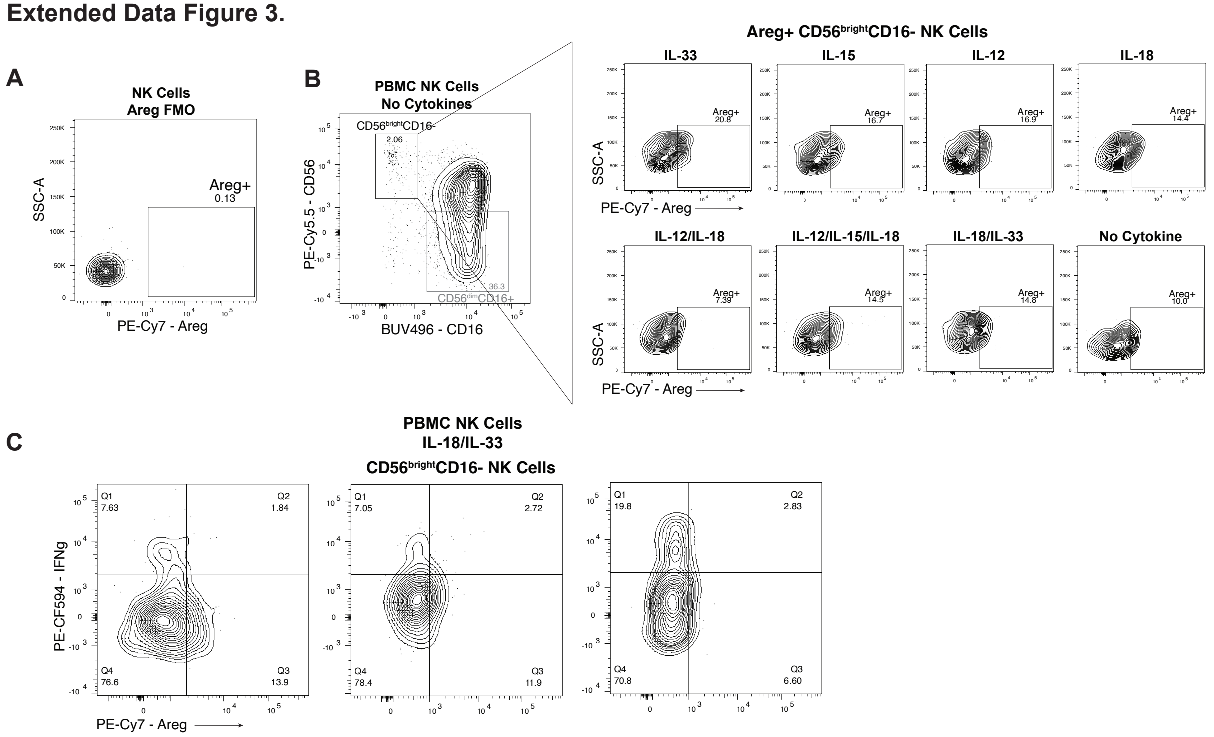


**Extended Data Figure 3.** Flow cytometry was performed on immune cells isolated from blood (PBMC) from healthy donors. Isolated PBMCs were stimulated with different combinations of cytokines (IL-12 10ng/mL, IL-15 50ng/mL, IL-18 10ng/mL, IL-33 50ng/mL) in complete RPMI media with 50ng/mL of IL-2 for 5 days at 37C, and the frequency of NK cells expressing Areg by CD56$^{bright}$ and CD56$^{dim}$ populations are graphed. **A**) Example flow plots of Areg FMO control demonstrating Areg- population. **B**) Example flow plots of NK cell CD56$^{bright}$CD16$^{-}$ populations expressing Areg from stimulated PBMCs. **C**) Representative flow plots from three donors showing PBMC NK cells expressing Areg and IFNg after IL-18 and IL-33 stimulation.

**Extended Data Figure 4.**

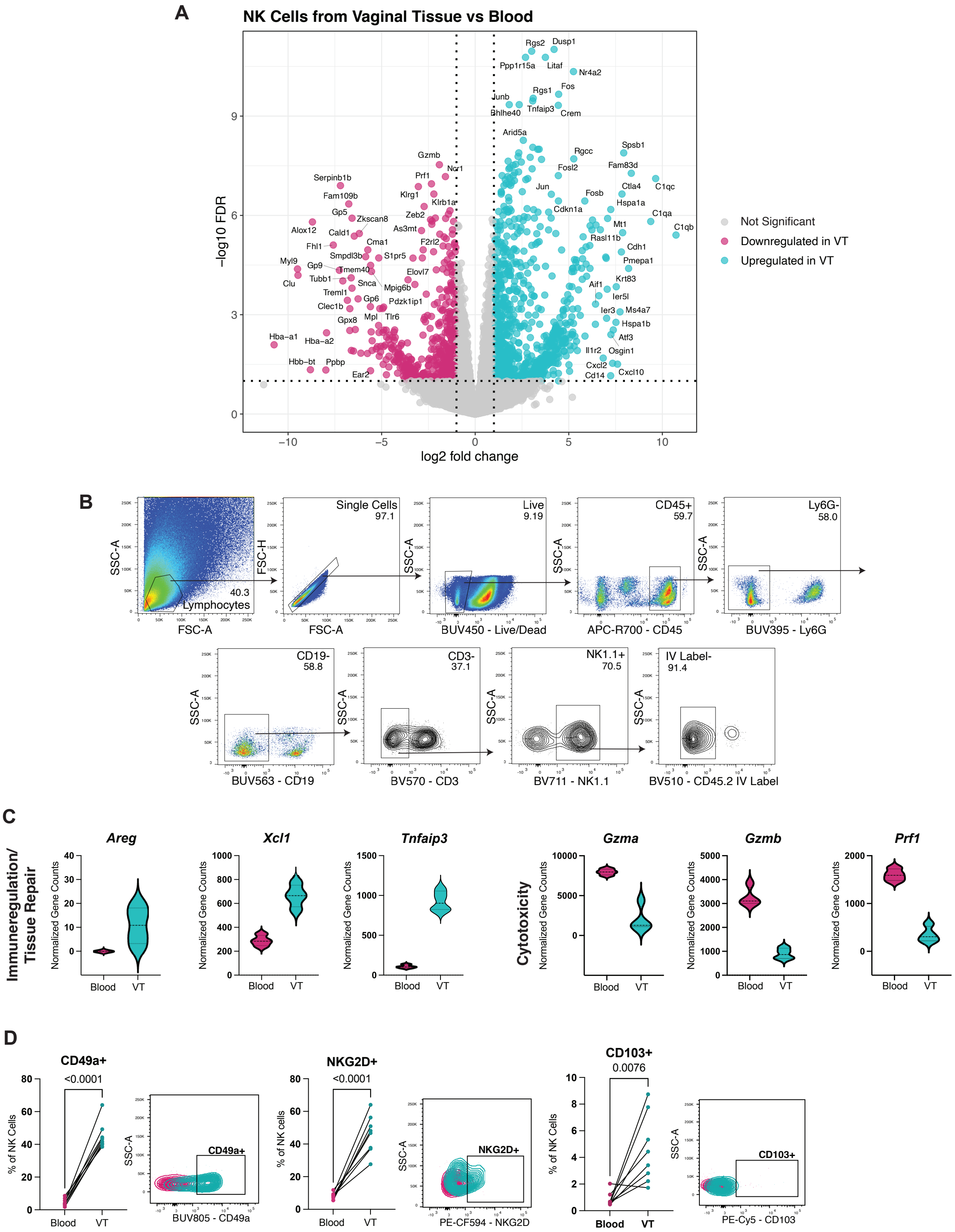


**Extended Data Figure 4.** Vaginal tissues and blood were harvested from uninfected C57BL/6 female mice. **A**) Volcano plot showing differentially expressed genes (DEG)

comparing NK cells isolated from blood to VT. **B**) Flow cytometry was performed, and NK cells were gated on Lymphocytes/SingleCells/Live/ CD45+/Ly6G-/CD19-/CD3-/NK1.1+/IVLabel-. **C**) Violin plots showing normalized gene counts for selected genes from NK cells isolated from blood (left) or VT (right). NK cells expressing markers of tissue repair and cytotoxicity. RNA-seq data show n=4 for blood and VT samples. **D**) Example flow plot and graphed frequencies of NK cells expressing markers of immune regulation and tissue residency (CD49a, NKG2D, CD103). All flow cytometry graphs show data for n=8 blood samples and VT samples. Data are combined from two independent experiments. Statistical analyses were performed using paired t test, with p values listed.

**Extended Data Figure 5.**

A

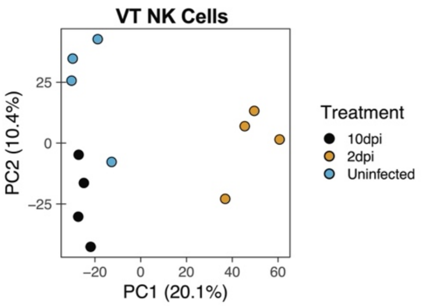


B

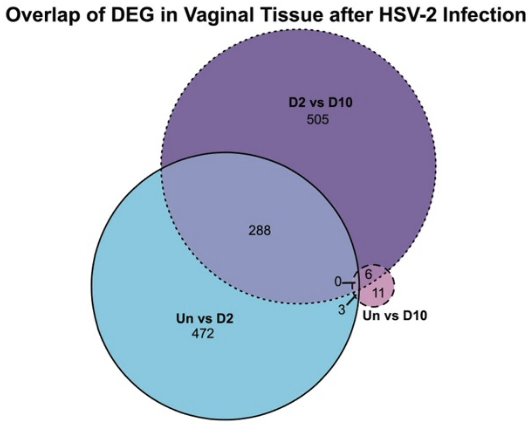


C

| Gene Name | Adj P Value | P Value | Log FC | Ave Expr |
|---|---|---|---|---|
| Ophn1 | 0.00926 | 1.289E-06 | 5.462408208 | -0.213485325 |
| H2-Eb1 | 0.00926 | 1.449E-06 | 1.741776086 | 8.826902849 |
| Cd74 | 0.01180 | 2.770E-06 | 1.378036486 | 10.91405611 |
| B4galnt4 | 0.01316 | 4.751E-06 | 5.105616582 | -0.372599423 |
| H2-Ab1 | 0.01316 | 5.193E-06 | 1.516305235 | 10.09766727 |
| Slco2b1 | 0.01316 | 6.179E-06 | 5.129204376 | -0.384972512 |
| Hepacam2 | 0.02071 | 1.134E-05 | 5.091493823 | -0.365524339 |
| Icos | 0.02216 | 1.387E-05 | 1.246313428 | 7.375259493 |
| Ly6c2 | 0.03314 | 2.334E-05 | 1.334682815 | 8.506677898 |
| Rasgrp3 | 0.03481 | 2.724E-05 | 4.890693872 | -0.47193456 |
| Tlr1 | 0.03640 | 3.133E-05 | 5.069943518 | -0.461963912 |
| Rgs1 | 0.04581 | 4.302E-05 | 0.529401943 | 10.85184785 |
| Gzmc | 0.06194 | 6.580E-05 | 1.892803905 | 7.68557416 |
| Themis | 0.06194 | 6.983E-05 | 5.815442207 | 1.777030002 |
| Gpr146 | 0.06194 | 7.270E-05 | 4.93798991 | -0.538904373 |
| Birc5 | 0.06252 | 7.827E-05 | 4.520578119 | -0.711305783 |
| Cxcl16 | 0.06366 | 8.469E-05 | 2.138139711 | 5.109679116 |
| Nr4a2 | 0.06624 | 9.587E-05 | -0.449660781 | 10.42381481 |
| Lifr | 0.06624 | 9.848E-05 | 4.690689902 | 0.294248618 |
| H2-Aa | 0.06924 | 1.100E-04 | 1.582778043 | 9.198374477 |
| Cd3d | 0.06924 | 1.144E-04 | 1.599834524 | 6.290883329 |
| Trim30b | 0.06924 | 1.192E-04 | -4.4844462 | -0.626049946 |
| Gramd3 | 0.07029 | 1.311E-04 | 1.573475529 | 7.55898765 |
| Il7r | 0.07029 | 1.320E-04 | 1.883157489 | 6.018575572 |
| Sqstm1 | 0.07437 | 1.455E-04 | -0.510683149 | 10.45422808 |
| Dclk2 | 0.07566 | 1.539E-04 | -4.242488899 | -0.649171185 |
| Slc40a1 | 0.09092 | 1.921E-04 | -4.536230687 | 0.200204492 |
| Trem3 | 0.09102 | 2.052E-04 | -4.297349232 | 0.019099338 |
| Lair1 | 0.09102 | 2.138E-04 | 1.242636569 | 5.827403983 |
| Mtus1 | 0.09102 | 2.219E-04 | 5.395755243 | 1.302002107 |
| Cd3e | 0.09102 | 2.271E-04 | 2.181785216 | 4.921864353 |
| Nr6a1 | 0.09102 | 2.307E-04 | -4.22870804 | -0.684238321 |
| Cd83 | 0.09102 | 2.409E-04 | 2.651986413 | 5.71948613 |
| Miip | 0.09102 | 2.421E-04 | -0.851064033 | 6.237538491 |
| H2-DMb1 | 0.09700 | 2.657E-04 | 1.349724063 | 5.664544116 |
| Sgcb | 0.09726 | 2.740E-04 | -2.10148966 | 4.318167303 |

**Extended Data Figure 5.** Female C56BL/6J mice were infected with 2x$10^6$ PFU HSV-2 TK- or left uninfected. Vaginal tissue (VT) and blood was harvested from uninfected mice, mice at 2dpi, and mice at 10dpi. 250 NK cells were sorted from each group gated on Live/CD45+/ Ly6G-/CD19-/CD3-/NK1.1+ and bulk RNA-seq was performed. **A**) PCA plot from bulk RNA-seq data on sorted NK cells demonstrating differential clustering of NK cells from uninfected mice (n=4), 2dpi (n=4), and 10dpi (n=4). **B**) Venn diagram showing the number of DEGs between each comparison and the number of overlapping DEGs between groups. **C**) List of DEGs comparing VT NK cells from uninfected mice to 10dpi with an adj p value cut off of 0.1 (n=4 for each group).

### Extended Data Figure 6.

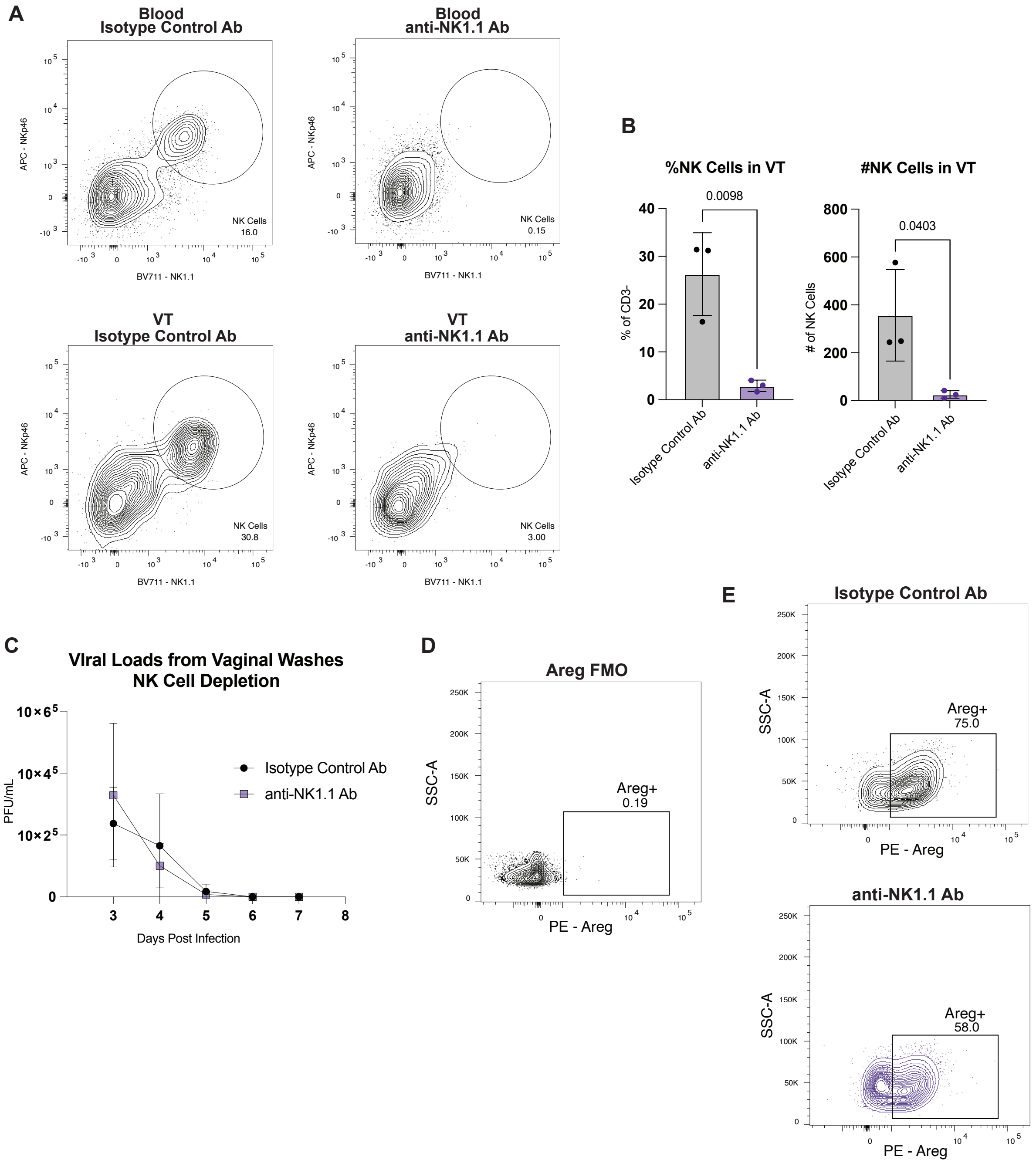

**Extended Data Figure 6.** Female C57BL/6 mice were infected with $2x10^6$pfu HSV-2 TK- and administered 200 µg of NK cell depleting antibody anti-NK1.1 or IgG2a isotype control antibody on days +4, +5, and +8 after infection. Vaginal tissues (VT) and blood from mice were harvested on day +10 after infection for flow cytometry. **A**) Representative flow plots from blood (top panels) or VT (lower panels) confirming NK cell depletion in mice given 200ug anti-NK1.1 Ab or isotype control Ab. Cells were gated on Live/CD45+/Ly6G-/CD19-/CD3-/NK1.1+/NKp46+. **B**) Graphs showing frequency and total number of NK cells in VT from anti-NK1.1 or isotype control groups. Data representative of two independent experiments. Graphs show mean with SD of n=3 mice and statistical analysis performed using unpaired t test, p values listed. **C**) Viral loads from vaginal washes taken from mice daily after infection with HSV-2 TK-. Viral titers were calculated using plaque assays day +3 post infection, before NK cell depletion and then daily until day +8 post infection when the virus was cleared (n=4 isotype control, n=4 anti-NK1.1). **D**) Areg FMO flow cytometry gating control. Cells gated on Live/CD45+. **E**) Representative flow plot demonstrating Areg staining on CD45+Live cells in the VT at 10dpi.
